# Molecular basis of polyspecific drug binding and transport by OCT1 and OCT2

**DOI:** 10.1101/2023.03.15.532610

**Authors:** Yang Suo, Nicholas J. Wright, Hugo Guterres, Justin G. Fedor, Kevin John Butay, Mario J. Borgnia, Wonpil Im, Seok-Yong Lee

## Abstract

A wide range of endogenous and xenobiotic organic ions require facilitated transport systems to cross the plasma membrane for their disposition^1, 2^. In mammals, organic cation transporter subtypes 1 and 2 (OCT1 and OCT2, also known as SLC22A1 and SLC22A2, respectively) are polyspecific transporters responsible for the uptake and clearance of structurally diverse cationic compounds in the liver and kidneys, respectively^3, 4^. Notably, it is well established that human OCT1 and OCT2 play central roles in the pharmacokinetics, pharmacodynamics, and drug-drug interactions (DDI) of many prescription medications, including metformin^5, 6^. Despite their importance, the basis of polyspecific cationic drug recognition and the alternating access mechanism for OCTs have remained a mystery. Here, we present four cryo-EM structures of apo, substrate-bound, and drug-bound OCT1 and OCT2 in outward-facing and outward-occluded states. Together with functional experiments, *in silico* docking, and molecular dynamics simulations, these structures uncover general principles of organic cation recognition by OCTs and illuminate unexpected features of the OCT alternating access mechanism. Our findings set the stage for a comprehensive structure-based understanding of OCT-mediated DDI, which will prove critical in the preclinical evaluation of emerging therapeutics.

## Main

OCTs are members of the solute carrier 22 (SLC22) transporter family. OCT subtype 1 (OCT1; SLC22A1) is highly expressed in the liver, whereas OCT2 (SLC22A2) is primarily expressed in the kidney^2^. OCT1 and OCT2 exhibit similar substrate specificity, transporting various endogenous cationic compounds such as thiamine, uremic solutes, and biogenic amines (e.g. epinephrine, serotonin, and dopamine)^7–10^. Notably, OCT1 and OCT2 respectively mediate the hepatic uptake and renal secretion of a wide range of cationic drugs, and play critical roles in drug disposition and response^11^. Case in point, the gold standard type II anti-diabetic drug metformin is principally taken up into the liver and kidneys by OCT1 and OCT2, respectively. Consequently, many genetic variants of *slc22a1* and *slc22a2* are associated with decreased metformin responses and altered pharmacokinetics^12–14^. Likewise, recent studies of genetic polymorphisms demonstrate the key role of OCT1 and OCT2 in the pharmacokinetics and pharmacodynamics of many drugs and controlled substances^11, 15–18^. There are currently well over 250 identified prescription drugs that are either substrates or inhibitors of OCT1 and OCT2, with a growing list that includes diphenhydramine (antihistamine), fluoxetine and imipramine (antidepressants), and imatinib (anticancer)^19–21^.

The polyspecificity of hOCT1 and hOCT2, and the fact that approximately 40% of prescription medicines are organic cations^22^, highlights their role in transporter mediated drug-drug interactions (DDI). This suggests broad implications on drug and clinical trial design, as DDI is a critical factor in clinical drug disposition, response, and toxicity. In fact, hOCT1 and hOCT2 have been implicated in multiple DDI instances^6, 21, 23, 24^. For example, the antihypertensive drug verapamil, which is an OCT1 inhibitor, was shown to decrease the glucose-lowering effect of metformin through DDI on hOCT1^25^. Because untested DDIs may introduce severe adverse effects on patients, the European Medicines Agency (EMA), the US Food and Drug Administration (FDA), and the International Transporter Consortium recommend *in vitro* testing of new therapeutics for potential interaction with hOCT1 and hOCT2^11, 26, 27^.

Over the past few decades, a wealth of functional studies has uncovered several key features of substrate recognition and drug interaction with hOCT1 and hOCT2^28–36^. However, the structural basis of substrate recognition, transport inhibition, DDI, and the transport mechanism of hOCT1 and hOCT2 remain elusive. The polyspecificity of hOCT1 and hOCT2 is in stark contrast with other SLC transporters, making it challenging to postulate a common binding mode and their transport mechanism in the absence of structural information. This ultimately hinders the development of more accurate methods to critically evaluate novel therapeutics for their interaction with hOCT1 and hOCT2 at the preclinical stage^4, 19^.

### Structure determination of OCT1 and OCT2

Wild type human OCT1 and OCT2 (WT hOCT1, WT hOCT2) express poorly in transiently transfected HEK293T cells, which prohibited biochemical optimization (Extended Data Fig. 1a). To enable structural studies, we turned to consensus mutagenesis^37, 38^. This approach resulted in two engineered OCT proteins, which we term OCT1_CS_ and OCT2_CS_ (see Methods for description of consensus construct design). Exhibiting sequence identities of 87% and 83% to WT, respectively (Extended Data Fig. 1b), both constructs express well in transiently transfected HEK293T cells and exhibit monodisperse behavior in fluorescence size exclusion chromatography (FSEC) analysis (Extended Data Fig. 1a). When expressed in *Xenopus laevis* oocytes, OCT1_CS_ mediates accumulation of tritiated 1-methyl-4-phenylpyridinium (^3^H-MPP^+^) to levels higher than WT hOCT1 (Extended Data Fig. 1c), while retaining intrinsic transport properties similar to WT. This is evidenced by a determined *K_t_* value of ∼50 μM for MPP^+^ transport by OCT1_CS_, which is consistent with previous reports for WT hOCT1 (Fig. 1a)^9, 39, 40^. Also, IC_50_ values for verapamil (VPM) or the antihistamine diphenhydramine (DPH) are similar for WT hOCT1 and OCT1_CS_, based on cold competition of [^3^H]-MPP^+^ uptake in oocytes (Fig.1b, c). Furthermore, the OCT1_CS_ *K_t_* value of ∼3 mM for metformin (Fig. 1d) is within range of previous reports for WT hOCT1 (1- 5 mM)^13, 41^. Finally, WT and OCT1_CS_ are functionally similar in cold-competition experiments against ^14^C-metformin transport with MPP^+^, VPM, DPH, and imatinib (IMB) (Fig 1e, Extended Data Fig. 1d). OCT2_CS_ is also functionally competent, exhibiting higher raw ^3^H-MPP^+^ uptake relative to WT hOCT2 with similar levels of block by racemic VPM (Fig. 1f, Extended Data Fig. 1e). Additionally, OCT2_CS_ does not mediate uptake of [^3^H]-VPM, consistent with VPM being an OCT2 inhibitor^2^, and exhibits only a modest stereoselectivity for VPM inhibition (Extended Data Fig. 1f,g).

**Figure 1.**
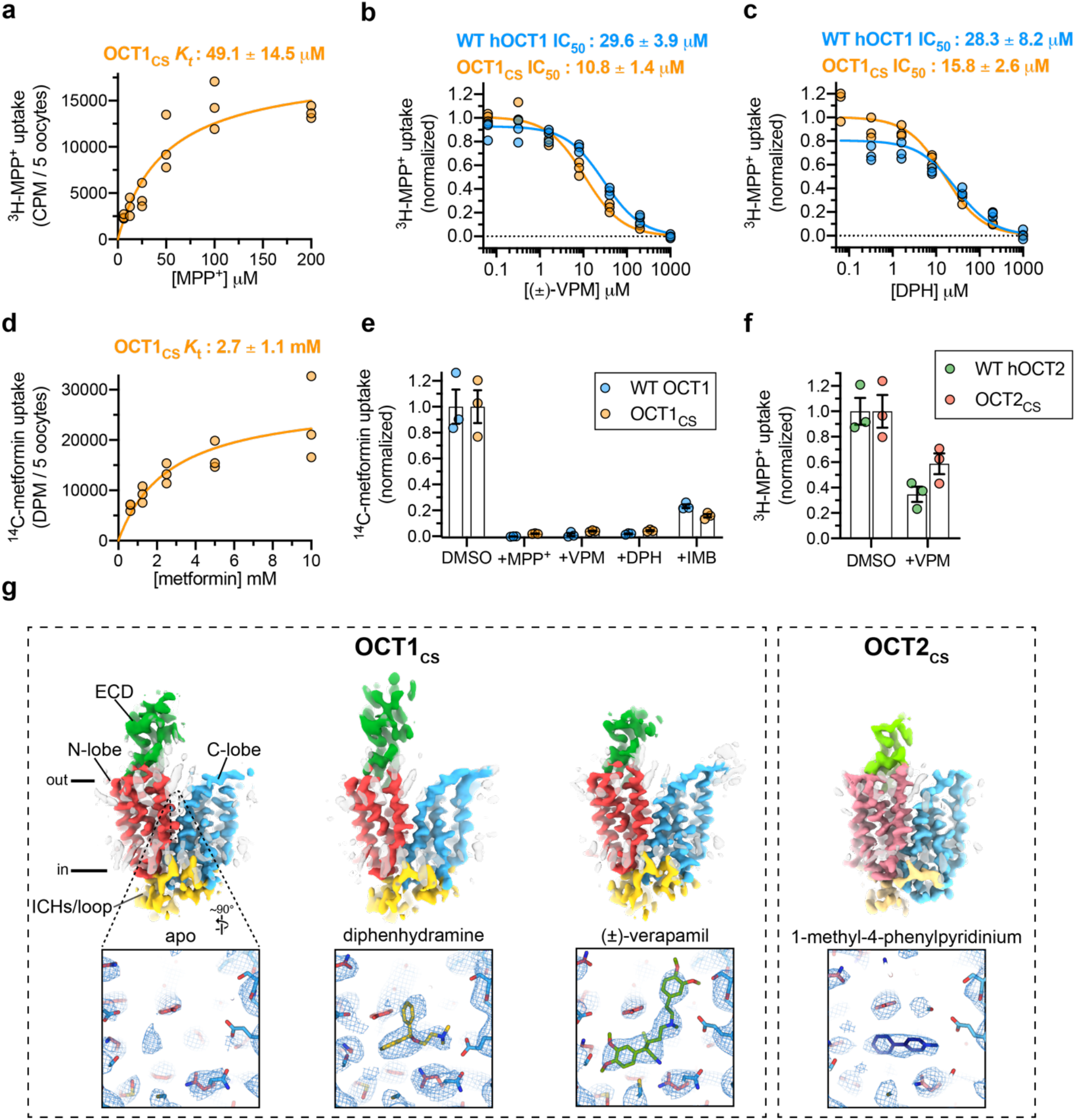
Cryo-EM structures of organic cation transporters 1 and 2. **a,** *K_t_* determination for ^3^H-MPP^+^ uptake mediated by *X. laevis* oocytes expressing OCT1_CS_ (30 minute uptake; *n*=3 individual biological replicates shown, *K_t_* ± s.e.m.) **b,** Cold-competition inhibition of WT hOCT1 or OCT1_CS_ mediated ^3^H-MPP^+^ uptake by (±)-VPM (30 minute uptake with 100 nM ^3^H-MPP^+^; *n*=3 individual biological replicates shown, IC_50_ ± s.e.m.). **c,** Cold-competition inhibition of WT hOCT1 or OCT1_CS_ mediated ^3^H-MPP^+^ uptake by DPH (30 minute uptake with 100 nM ^3^H-MPP^+^; *n*=3 individual biological replicates shown, IC_50_ ± s.e.m.). **d,** *K_t_* determination for ^14^C-metformin uptake mediated by OCT1_CS_ (*n*=3 individual biological replicates shown, *K_t_* ± s.e.m.). **e,** Single concentration-point cold-competition block of ^14^C-metformin uptake (83.3 μM) by WT hOCT1 or OCT1_CS_ with 1 mM cold MPP^+^, VPM, or DPH, or 0.231 mM cold IMB (*n*=3 individual biological replicates shown, mean ± s.e.m.; unnormalized water, WT and OCT1_CS_ injected controls shown in Extended Data Fig. 1c for reference). **f,** WT hOCT2 or OCT2_CS_ mediated ^3^H-MPP^+^ uptake (1 hour uptake with 100 nM ^3^H-MPP^+^; *n*=3 individual biological replicates shown, mean ± s.e.m.; unnormalized values for water, WT and OCT2_CS_ injected controls shown in Extended Data Fig. 1c for reference). **g,** Cryo-EM reconstructions of apo-OCT1_CS_, DPH- OCT1_CS_, VPM-OCT1_CS,_ or MPP-OCT2_CS_ (top), with cryo-EM densities of the central cavity shown at bottom (map thresholds are set at 0.45, 0.25, 0.30, or 0.25 for apo-OCT1_CS_, DPH-OCT1_CS_, VPML-OCT1_CS_ , or MPP^+^-OCT2_CS_ ligand densities, respectively).

Owing to the enhanced expression level and biochemical stability of OCT1_CS_ and OCT2_CS_, (Extended Data Fig. 1a, e), we then solved three cryo-EM structures of OCT1_CS_: in the absence of added ligand to 3.57 Å resolution (apo-OCT1_CS_), with (±)-VPM bound to 3.45 Å resolution (VPM- OCT1_CS_), and with DPH bound to 3.77 Å resolution (DPH-OCT1_CS_; Fig. 1g, Extended Data Figs. 1f-g, and 2-3, Extended Data Table 1). We also solved one structure of OCT2_CS_: with MPP^+^ bound to 3.61 Å resolution (Fig. 1g and Extended Data Figs. 2-3, Extended Data Table 1). The local resolutions for the ligand/ligand binding regions for apo-OCT1cs, VPM-OCT1_CS_, DPH-OCT1_CS_, and MPP^+^-OCT2_CS_ are ∼3.3, ∼3.1, ∼3.5, and ∼3.4, respectively (Fig. 1g and Extended Data Figs. 2-3).

**Figure 2.**
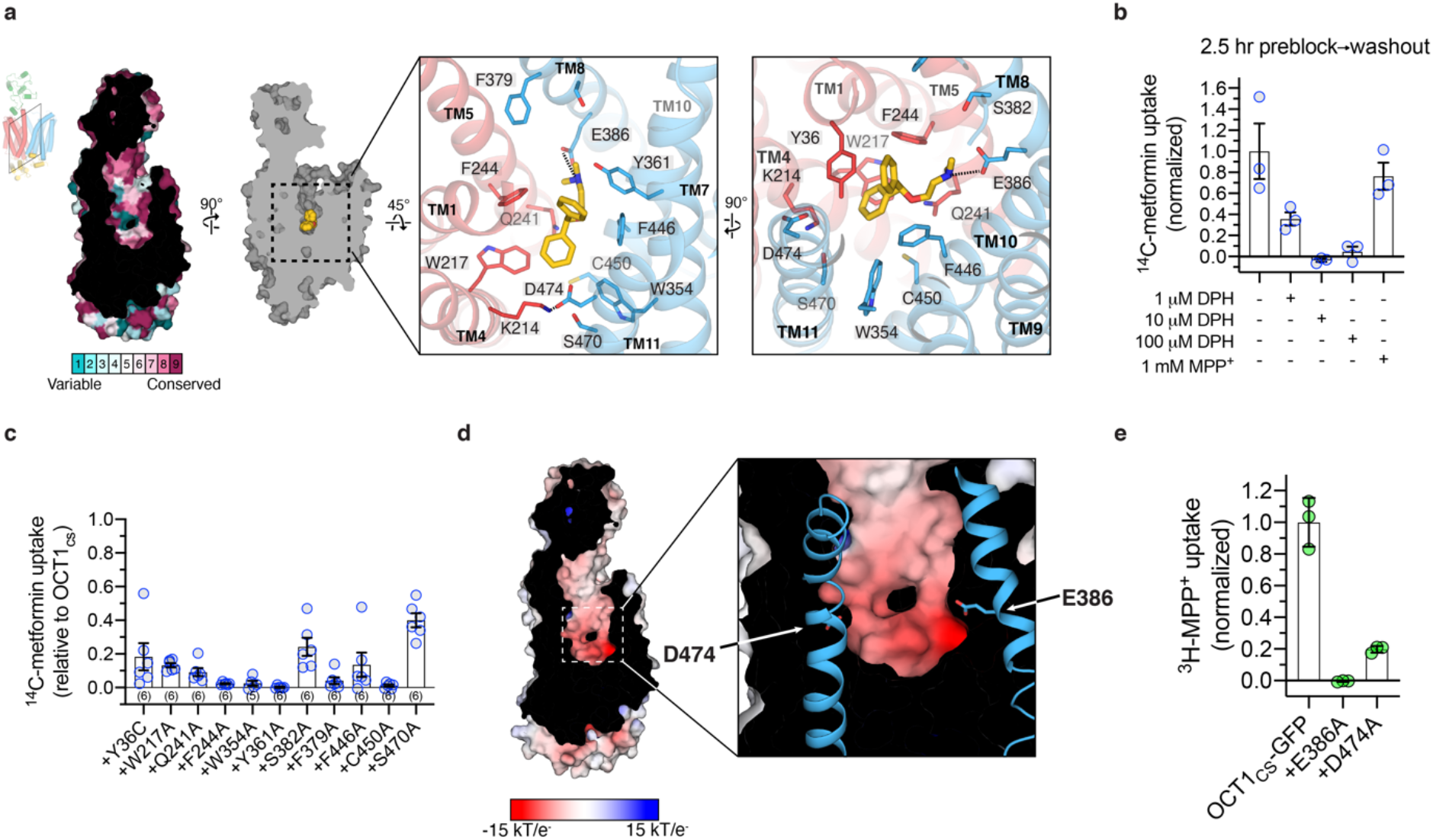
Diphenhydramine recognition by OCT1. **a**, ConSurf^53^ analysis of the OCT1_CS_ central cavity (left). Residue Y36 in the central cavity shows high variability across OCT1 orthologs in the multiple sequence alignment. Detailed DPH-OCT1 interactions in the binding cavity (right), highlighting interacting residues. **b**, Cold competition block of ^14^C-metformin uptake mediated by OCT1_CS_ (10 μM in 30 minutes) after 2.5-hour pre- treatment at the noted concentration, followed by rapid and extensive oocyte washing in ligand- free buffer (see Methods for details). **c,** Functional evaluation of mutants in the OCT1_CS_ background (accumulation of 10 μM ^14^C-metformin in 60 min into mutant-expressing oocytes; *n* individual biological replicates shown as indicated in parenthesis, mean ± s.e.m.). **d**, APBS^54^ surface electrostatic calculation of the OCT1_CS_ central cavity (see Methods). **e,** Uptake of ^3^H- MPP^+^ by OCT1_CS_-GFP or mutants in the OCT1_CS_-GFP background (accumulation of 100 nM ^3^H- MPP^+^ in 60 min into mutant-expressing oocytes; *n*=3 with individual biological replicates shown along with mean ± s.e.m.)

**Figure 3.**
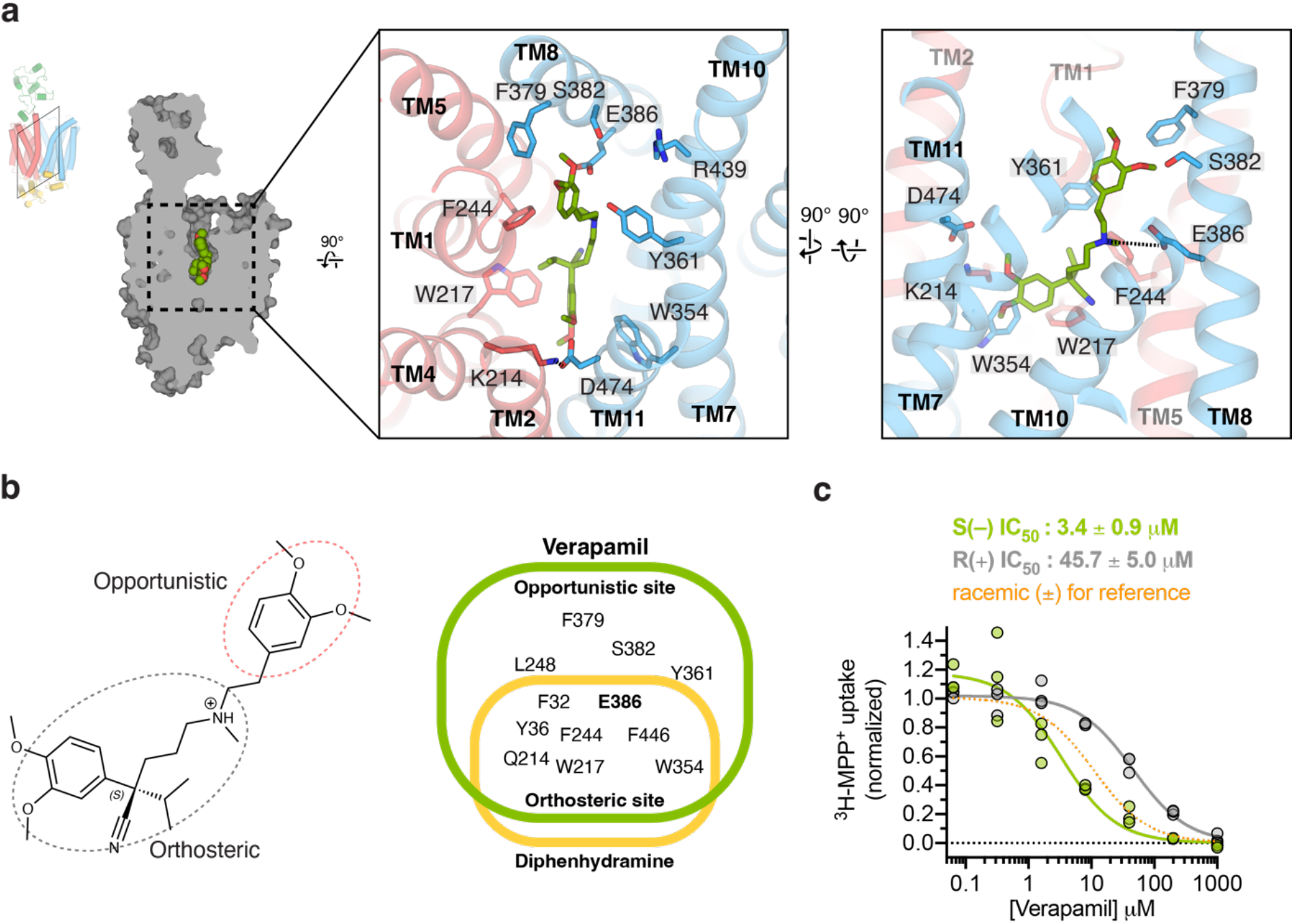
Verapamil recognition by OCT1. **a**, Detailed VPM-OCT1 interactions in the binding cavity, highlighting interacting residues. **b**, Orthosteric and allosteric moieties of VPM (left), with shared and distinct interacting residues between VPM and DPH (right). **c**, Enantiospecific recognition of VPM by OCT1_CS_, as shown by IC_50_ measurements of S(–) or R(+) VPM against OCT1_CS_ mediated ^3^H-MPP^+^ uptake activity (30 minute uptake with 100 nM ^3^H-MPP^+^; *n*=3 individual biological replicates shown, IC_50_ ± s.e.m.).

The overall OCT fold can be divided into three parts – an extracellular domain (ECD), a transmembrane domain consisting of 12 transmembrane (TM) helices and an intracellular helical (ICH) bundle comprised of four short helices (Fig. 1g, Extended Data Fig. 1h). The 12 TM helices form a 6+6 pseudosymmetrical arrangement with TMs 1-6 comprising the N-terminal lobe, and TMs 7-12 the C-lobe. The reconstructions obtained in the presence of ligand feature well-defined densities in the central cavity between the N- and C- domains, while the apo-OCT1_CS_ reconstruction lacks such density (Fig 1g).

The interface between the N- and C-lobes of OCTs form a highly conserved cavity in which substrates bind (Fig. 2a, Supplemental Fig. 1). All three OCT1_CS_ reconstructions, apo-OCT1_CS_, DPH-OCT1_CS,_ and VPM-OCT1_CS_, adopt an apparent outward-facing open conformation, as the opening at the extracellular side is large enough to readily accommodate substrate entry. A feature unique to OCT1 is an extended extracellular domain (ECD) located between TMs 1 and 2. The ∼90 residue ECD forms a cap-like structure that sits atop the N-lobe and interacts with the TM3- TM4 and TM5-TM6 loops. Compared to the TMs, the ECD is sub-optimally resolved due to its relative flexibility (Fig. 1g).

### Diphenhydramine binding to OCT1

The robust cryo-EM density and molecular dynamics simulations of two possible binding poses allowed us to assign the DPH molecule in DPH-OCT1_CS_ without ambiguity (Extended Data Fig. 4a,b). The DPH molecule is stabilized by several hydrophobic residues (Fig. 2a), in particular W217 (TM4) and F244 (TM5) on the N-lobe, as well as W354 (TM7) and F446 (TM10) on the C-lobe (F446 is isoleucine in hOCT1). These four residues form opposing “walls” of the binding pocket, with the only two acidic residues within the cavity, E386 (TM8) and D474 (TM11), defining the other two sides (Fig 2a). The positively charged dimethylethanamine group interacts exclusively with E386, while D474 (TM11) forms a charge-pair with neighboring K214 (TM4) (Fig. 2a). Y36 (TM1) and Y361 (TM7) line the cavity above the plane of the four hydrophobic residues. Typically, MFS transporters bind ligand with residues on TMs 1, 4, 7, 10 (known as A helices)^42, 43^, however OCT1 also recruits TMs 5 and 8 (B helices) to bind DPH. Only two residues differ in the ligand binding cavity between OCT1_CS_ and hOCT1, Y36 (C36 in hOCT1) and F446 (I446 in hOCT1) (Fig. 2a and Extended Data. Fig. 1b).

**Figure 4.**
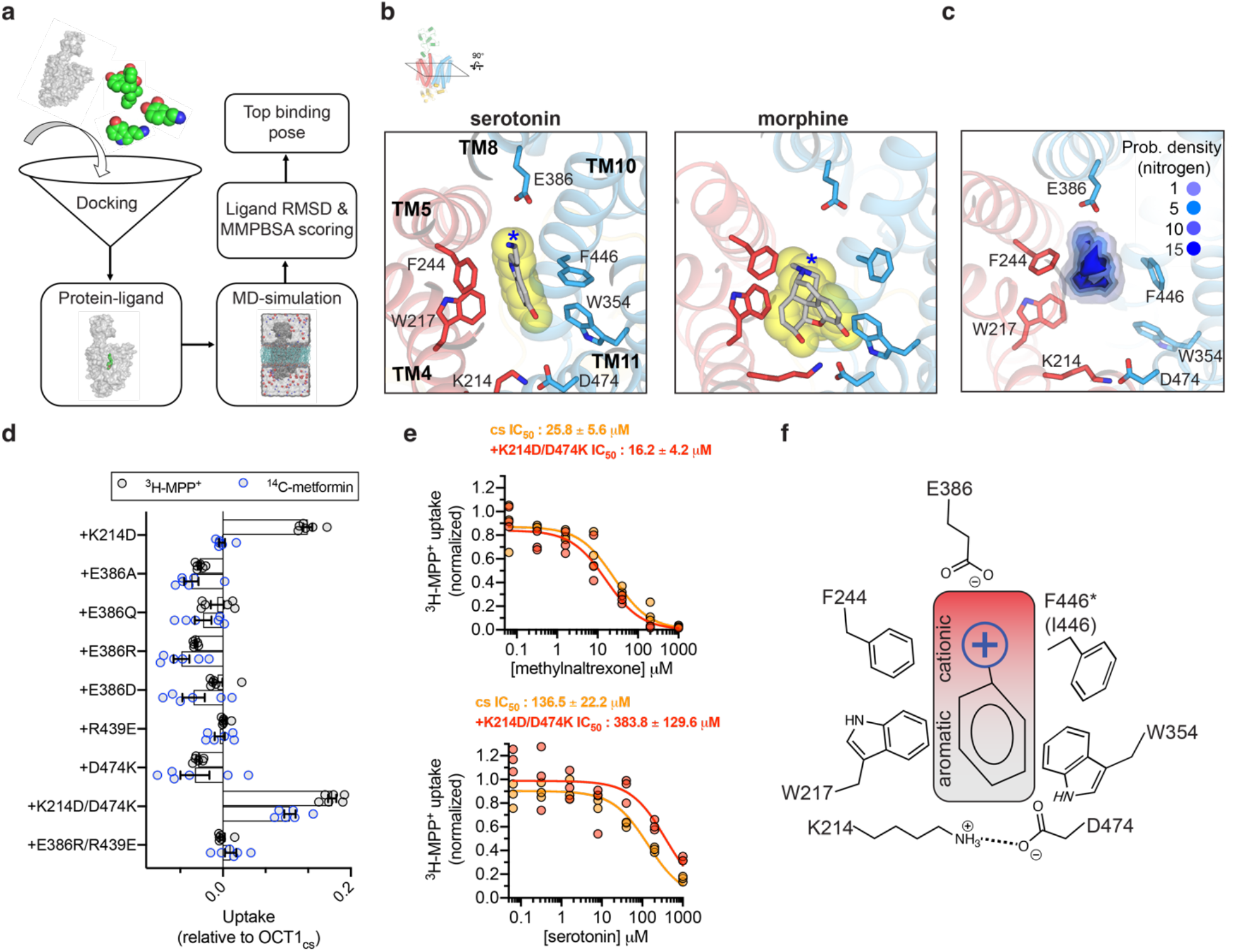
General principles of organic cation recognition by OCT1. **a,** Scheme for docking-MD predictions of drug binding poses. **b,** Final MD frames of top scored binding poses for two representative drugs. **c,** Probability density for basic-nitrogen atom positions in the ten interrogated drugs, from the final MD frame of top scored binding poses (threshold value are arbitrary) **d,** Uptake measurements for charged position mutants in the OCT1_CS_ background (either 10 μM ^14^C-metformin or 10 nM ^3^H-MPP^+^ uptake in 60 minutes; *n*=6 individual biological replicates shown with mean ± s.e.m.). **e,** Inhibition of OCT1_CS_ or charge-swap double mutant (OCT1_cs_+K214D/D474K) by methylnaltrexone (top panel) or serotonin (bottom panel; *n*=3 individual biological replicates shown with IC_50_ ± s.e.m.). **f,** A general model for organic cation recognition by outward-facing OCT1.

It is not well established whether DPH is a transported substrate or inhibitor of OCT1. The quantity of radioactive DPH required for uptake experiments prohibited direct testing for OCT1_CS_ uptake of DPH, so we instead pursued cold ligand wash-out experiments. Unlike the OCT1 substrate MPP^+^, DPH exhibits apparently slow off-rate kinetics since substantial residual block of [^14^C]- metformin uptake remained after washing out external cold DPH (Fig. 2b). This data suggests DPH is a non-transported inhibitor of OCT1_CS_. Therefore, for further functional interrogation of OCT1_CS,_ [^14^C]-metformin uptake measurements were performed on alanine mutants of residues lining the central cavity (Fig 2c). We found that the aromatic and aliphatic residues interacting with DPH are also critical for metformin transport, as their alanine mutants show substantial reductions in transport activity. Their expression was verified by transfection of HEK293T cells (Extended Data Fig. 1d).

Electrostatic surface potential calculations show that the central cavity is anionic, with E386 appearing to make the greatest contribution (Fig. 2d). While previous studies have implicated D474 (numbering consistent with hOCT1) as being critical for cation binding and translocation in OCT1^29, 32, 44^, the role of E386 has never been interrogated to the best of our knowledge. We measured MPP^+^ uptake activity of E386A and D474A in the OCT1_CS_-GFP background, and while D474A retains ∼20% activity, E386A is dead (Fig. 2e). The oocyte surface expression of these constructs was confirmed by confocal microscopy (Extended Data Fig. 5). Our structural and functional observations reveal the importance of E386 in cation drug recognition by OCT1 (Fig 2a,d,e).

**Fig. 5.**
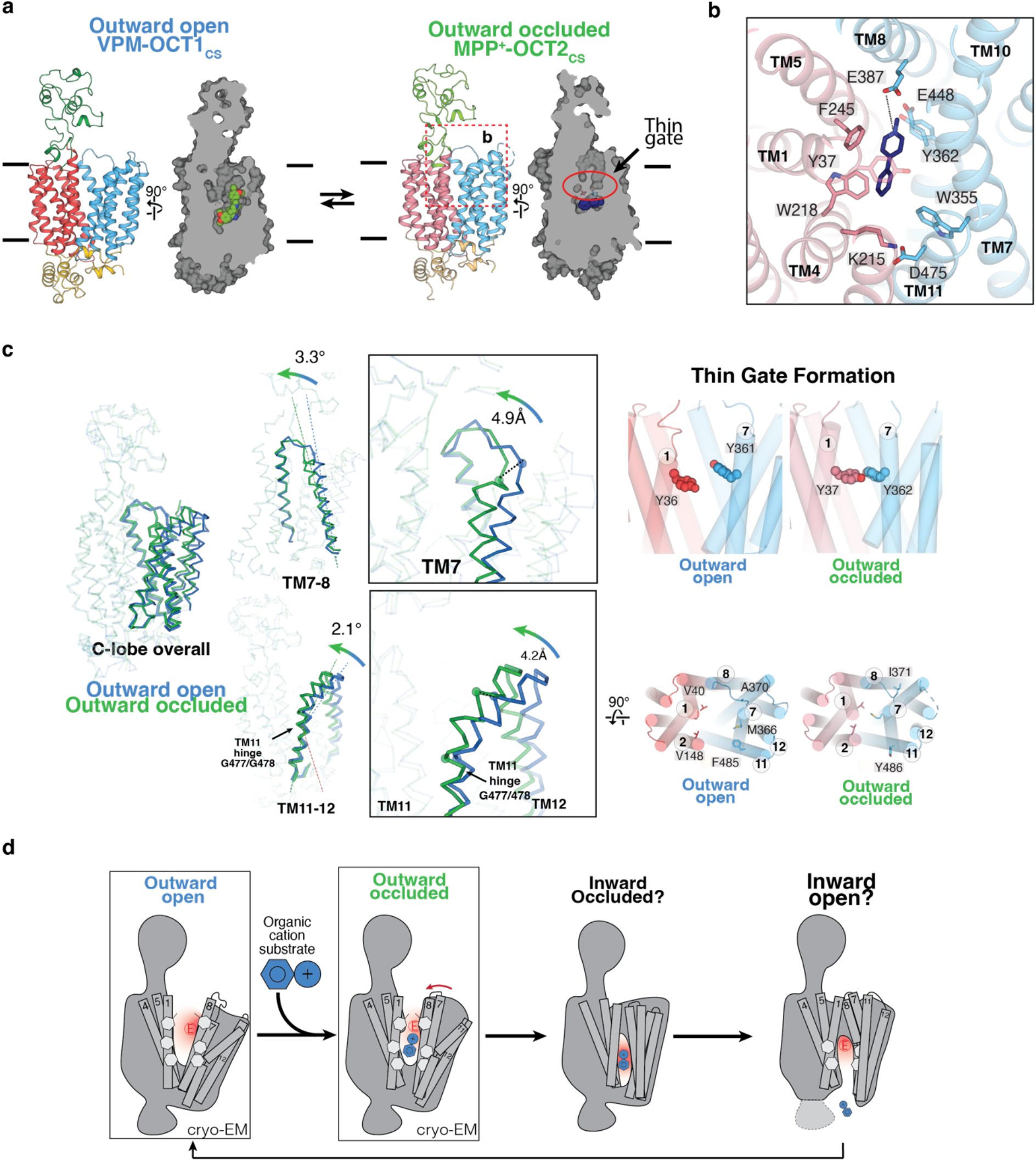
Extracellular gate closure in OCTs. **a**, Overview of the two distinct OCT conformations: outward open (VPM-OCT1_CS_) and outward occluded (MPP^+^-OCT2_CS_) **b,** Ligand binding pose for MPP^+^-OCT2_CS_, showing MPP^+^ (navy), and interacting residues of OCT2. N- and C- domains are colored blue and pink, respectively. **c,** Structural comparisons among the three observed conformations. Left, overall C-domain changes as well as TM7-8, TM11-12 conformational changes. Right, the conformational changes result in the formation of the thin and thick gates. **d,** Proposed OCT alternate access transport mechanism based on structural observations.

### OCT1 inhibition by verapamil

VPM is a well-established OCT1 inhibitor that creates a DDI with metformin via its inhibition of OCT1^25^. The high quality cryo-EM density for VPM, together with all-atom MD simulations of two possible binding poses, allowed unambiguous placement of a VPM molecule in the central cavity of OCT1_CS_ (Fig. 1g and Extended Data Fig. 4c,d). The drug moiety consisting of dimethoxyphenyl, isopropyl, and pentanenitrite groups is analogous to the diphenylmethoxy group of DPH and resides in the hydrophobic portion of the central cavity formed by the plane of four residues W217, F244, W354 and F446 (Fig 3a). Notably, the cationic tertiary amine group of verapamil forms a salt bridge with E386, as for DPH. Superposition of apo, DPH-, and VPM- bound OCT1 structures show rearrangements of Y36 in TM1 (cysteine in hOCT1 and tyrosine in rat OCT1) upon binding of different ligands, but only minor deviation for the hydrophobic plane residues (W217, F244, W354, and F446) and E386 (Extended Data Fig. 6).

The striking similarity of binding modes between VPM and DPH, and that E386A is devoid of MPP^+^ uptake activity despite its surface expression, led us to hypothesize the general roles of the acidic residue E386 in charge stabilization and aromatic/aliphatic residues W217, F244, W354 and F446 in hydrophobic packing of OCT1-bound compounds. We term this binding site as the orthosteric site. VPM also possesses another 3,4-dimethoxyphenyl group that extends toward the extracellular side of OCT1 (Fig 3b). The 3-methoxy group hydrogen bonds with S382 in TM8, and the phenyl group interacts with Y361 in TM7. This site, which we term the opportunistic site, is distinct from the orthosteric site as only larger substrates and/or inhibitors would presumably occupy it. Because the opportunistic site is proximal to the extracellular side of the transporter, binding of moieties to this site likely prevents the conversion from outward-facing open to outward-facing occluded, which may explain the inhibition of OCT1 by VPM. Similar modes of inhibition have been observed in other MFS transporters^38, 45, 46^.

In addition, it is worth noting that clinically utilized VPM is a racemic mixture, which we used for our cryo-EM sample preparation. There are many studies describing stereoselectivity-dependent target activity, pharmacokinetics, and pharmacodynamics of VPM^47–49^. The high quality ligand density in our cryo-EM reconstruction (Fig. 1g, Extended Data Fig. 4e) as well as the chemical environment of the orthosteric binding site supports binding of the S(–)-VPM enantiomer. In stereoselectivity experiments we found that S(–)-verapamil is ∼10 times more potent than R(+)- VPM in inhibiting ^3^H-MPP^+^ uptake in oocytes expressing our consensus construct (Fig. 3c). Thus indicating OCT1 preferably binds S(–)-VPM. Consistent with this observation, it was reported that the hepatic bioavailability of S(–)-VPM is lower than R(+)-VPM due to stereoselective first-pass metabolism^48, 49^.

### Insights into polyspecific organic cation recognition by OCT1

OCT1-interacting drugs are structurally diverse with only vague similarities (i.e. presence of a basic nitrogen connected to additional aromatic/aliphatic moieties), so we sought to utilize the OCT1_CS_ structures reported here for *in silico* binding mode prediction studies. To increase the prediction accuracy, our *in silico* binding prediction method is comprised of three stages. First, ligand binding mode is predicted using our holo structures. Second, stability is ascertained by all-atom MD simulations of the top ten predicted poses. Third, molecular mechanics with Poisson-Boltzmann electrostatic continuum solvation and surface area (MMPBSA) free energy calculations of stable binding poses (ligand root-mean-square deviation (R.M.S.D) < 3 Å) allow selection of a single top binding pose per ligand^50, 51^(Fig. 4a). Using this strategy, we were able to predict binding poses of VPM and DPH similar to those observed in the cryo-EM structures (Extended Data Fig. 7). We then predicted the binding modes of a small, diverse subset of known OCT1 ligands (Fig. 4b, Extended Data Fig. 7, Extended Data Table 2).

In most cases, the top scored pose from our docking-MD-free energy calculation strategy predicts that the aromatic/aliphatic groups pack against the aromatic residues proximal to the K214-D474 charge pair (Extended Data Fig. 7). Universally, however, the basic nitrogen of the drug is offset towards E386 or equidistant from D474 and E386 (Fig. 4c, Extended Data Fig. 4f, 7). Further, In the top poses of serotonin, mescaline, methylnaltrexone, imipramine, and MPP^+^, the basic nitrogen atom is closer to E386 than D474 in the predicted pose. To further probe the role of these two acidic residues, we systematically mutated both positions in the OCT1_CS_ background and performed radiotracer uptake assays in oocytes for both [^14^C]-metformin and [^3^H]-MPP^+^ (Fig. 4d). Notably, E386 is intolerant to mutation, as no E386 mutant yielded measurable signals in the OCT1_CS_ background (Fig. 4d). It is worthwhile to reiterate our finding that D474A retains some ^3^H-MPP^+^ uptake activity while E386A is nonfunctional, with both exhibiting comparable levels of surface expression in oocytes (Fig. 2e, Extended Data Fig. 5). Consistent with this finding, previous studies have also showed that substitutions at D474 are still functional in rat OCT1 (D475 in rat)^32, 52^. To probe the charge-pair between K214-D474, we assessed the charge-swap double mutant (D474K/K214D) in the OCT1_CS_ background and found that it partially rescues the loss of activity of D474K for both ^14^C-metformin and ^3^H-MPP^+^ uptake activity in oocytes (Fig. 4d). This contrasts with the E386R/R439E double mutant (∼9 Å apart in the VPM structure), which could not restore the activity of E386R. Furthermore, the D474K/K214D charge-swap mutant exhibited similar IC_50_ values for methylnaltrexone and serotonin (representative large and small OCT1 substrates, respectively), relative to OCT1_CS_ (Fig. 4e). This data further validates a charge pair between D474 and K214, while also suggesting there is low stringency for the precise positioning of the acidic residue at this side of the central cavity. Therefore, it is reasonable to suggest that D474 does not form a conserved, direct interaction with cationic substrates of OCT1 in the outward-facing state but rather helps tune the cavity electrostatics.

Therefore, drug recognition by OCT1 in the outward conformation involves the acidic residue E386 and aromatic/hydrophobic positions (217, 244, 354, 446) (Fig 4f), all of which provide the appropriate chemical environment capable of accommodating a wide range of cationic substrates. Like the needle of a compass, the cationic moiety of the drug orients in the OCT1 cavity toward E386. Our model is consistent with recent studies that identify high lipophilicity and a cationic charge as the main features required for drug binding to OCT1^19, 36^, features complementary to the binding site revealed by our structures. Multiple functional studies have suggested that OCT1 contains multiple binding sites that are either overlapping or allosteric^2, 19^. Our structural and functional studies demonstrate a core binding site of OCT1 in the outward state with the ability to accommodate extra moieties within an opportunistic site outside the orthosteric site.

### MPP^+^-bound OCT2 adopts the outward-facing occluded state

Unlike DPH and VPM, which inhibit transport, MPP^+^ is a well-established substrate of all OCT subtypes (Fig. 1a,1f, and^2, 44^). Notably, compared to the outward-facing open OCT1_CS_ structures the MPP^+^-OCT2_CS_ structure is more compact, adopting an outward-facing occluded conformation, (Fig. 5a). The high conservation is apparent in the central cavity between the two subtypes (Fig. 5b and Supplemental Fig. 1). MPP^+^ occupies space within the OCT2_CS_ central cavity that is analogous to the OCT1 orthosteric site. The clear cryo-EM density and limited number of possible ligand poses allowed us to model MPP^+^ confidently (Fig. 1g). While the 4-phenyl group closely interacts with OCT2 residues W218, F245, W355, and F447 (analogous to positions 217, 244, 354, and 446 in hOCT1), the 1-methylpyridinium group points towards E387 (E386 in hOCT1), with the charged nitrogen ∼4.8 Å from this acidic residue (Fig. 5a,b). OCT2 features an additional cavity lining acidic residue compared to OCT1, E448 (Q447 in hOCT1), which is also ∼5 Å from the charged nitrogen of MPP^+^ (Fig. 5b). Interestingly, the MPP^+^ binding pose observed here is consistent with what is predicted by *in silico* docking with OCT1 (Extended Data Fig. 7). Extracellular egress of MPP^+^ is blocked by Y37 (TM1) and Y362 (TM7; Fig. 5c) which form the thin gate in the outward-occluded state of MPP^+^-OCT2_CS_. Consistent with this observation, thin gate formation was obstructed in the VPM and DPH bound outward-open states of OCT1_CS_ due to their interactions with Y36 or Y361 (Fig. 2a, 3a, 5c and Extended Data Fig. 6).

### Insights into the OCT alternating access mechanism

Our structures of OCT1 and OCT2 in outward-open and outward-occluded conformational states yield unexpected insights into the alternating access mechanism of OCTs. The conformational changes from the outward-open to the outward-occluded states involve many local fold changes of both lobes. Specifically, TM7 rotates to form the extracellular thin gate, with TM11 forming a “latch” that clamps over TM7 during this transition, with helical movements occurring about a hinge point located at OCT1 G447 (G448 in OCT2) (Fig. 5c). A previous voltage-clamp fluorometry study implicated TM11 movements with MPP^+^ binding to rat OCT1^33^.

In addition to TM7, TM8, and TM11, local changes of TM2 are apparent. The interactions between TM2-3 and TM4, TM11, ICH2, and ICH3 in the outward facing structures stabilize the outward conformation (Extended Data Fig. 8c). TM2 which packs against TM11 is slightly rotated and offset upon thin-gate formation (Extended Data Fig. 8d). Considering the fact gating interactions mediated by TM2-3 are common in MFS transports^43^, it is possible differential interactions with ICH2-4 would be involved in transition between conformational states (Extended Data Fig. 8c).

## Discussion

Altogether, our cryo-EM structures, *in silico* drug docking-MD-free energy calculations, and functional experiments shed considerable insight into important features of both ligand recognition and the transport mechanism exhibited by OCTs. First, we discovered a shared motif of ligand recognition amongst the chemically diverse substrates of OCT1, in the context of the OCT1 outward-facing conformation. Considering the high tissue expression of OCT1 in the liver, this transporter conformation is relevant to first-pass metabolism of xenobiotics - where outward-facing OCT1 is poised to accept cationic drugs from the sinusoid. Second, with our structures we can infer alternate access rearrangements for OCTs. Local rearrangements of the B-helices (TM2, 5, 8, 11) in the MFS-fold are not typically associated with substrate binding and gating^43^. However, we observed the unexpected involvement of B-helices in ligand binding (TM5 and TM8) and conformational change (TM2, TM11), which to our knowledge is unprecedented. Previous functional data is consistent with the movement of TM11^33^, so it is tempting to speculate that the substantial local fold changes present at B helices between different conformational states facilitating greater plasticity in the substrate binding site during the transport cycle, allowing OCTs to translocate a wide range of cationic compounds (Fig. 5d).

It is important to reiterate that strategies for predicting the potential of new molecular entities for unwanted DDI are a critical aspect of therapeutic development^4^. The data presented here, including the *in silico* drug binding workflow utilized, could greatly accelerate drug development efforts. In total, our work sets the stage for structure-informed prediction of drug interactions with these two pharmacologically important polyspecific transporters at the preclinical stage.

## Materials and Methods

### Consensus mutagenesis design

Consensus constructs were designed in a similar manner to what has been previously reported in ^37^, with the following modifications. First, PSI-BLAST using WT hOCT1 or hOCT2 as the input (UniProt ID O15245) was performed to identify 250 OCT1 or OCT2 sequence hits from the UniProt database (nr90 - 90% similarity cut-off to reduce redundancy)^55^. To focus the sequence list to specific subtypes only, is was manually curated to select the top hits scored by sequence percentage identical to either subtype that were also annotated in the database as the particular subtype of interest. The remaining 58 sequences for OCT1 or 121 sequences for OCT2 were subjected to sequence alignment in MAFFT^56^. The consensus sequence was then extracted in JalView^57^, and aligned to the WT sequence in MAFFT. Sequence elements present in the WT sequence but not the consensus sequence (gaps in alignment present in loops and areas of low conservation) were then removed and replaced with WT sequence elements. The final constructs feature sequence registers consistent with WT.

### Oocyte radiotracer uptake assays

^14^C-metformin (115 Ci/mmol) was purchased from Moravek, and ^3^H-MPP^+^ (80 Ci/mmol) was purchased from American Radiolabeled Chemicals. Uptake assays were performed similarly to a previous report^58^. Injections of 30 ng cRNA were performed, with 2-4 day expression at 17°C. Specific radioactivities of 0.06 and 5 Ci/mmol were used for ^3^H-MPP^+^ and ^14^C-metformin for *K_t_* measurements shown in Fig. 1a and Fig. 1d, respectively. Full specific radioactivities were used for mutant uptake assessments in Fig. 1f, Fig. 2c, Fig. 2e, Fig. 4d. For IC_50_ experiments, specific radioactivities of 80 Ci/mmol were used for all constructs, except for OCT1_CS_ for which a specific radioactivity of 8 Ci/mmol was used. Water injected oocytes were used for background correction.

### Oocyte Fluorescence Microscopy

The method for fluorescence microscopy of oocytes was adopted from Löbel et al.^59^ with minor modifications. Oocytes were injected with either water or 30 ng of cRNA with protein expression occurring over 2 days at 17°C. Oocytes were harvested, washed twice with PBS, then stained with 0.05 mg ml^-1^ CF633-conjugated wheat germ agglutinin (Biotium) in PBS for 5 min at RT. Oocytes were then washed with PBS. CF633 and eGFP fluorescence were measured using a Leica SP8 upright confocal microscope equipped with a 10× objective lens, using He-Ne (633 nm) and Argon (488 nm) lasers, for CF633 and eGFP, respectively.

### OCT1/2 Protein expression and purification

Full-length consensus OCT1 and OCT2 sequences were codon-optimized for *Homo sapiens* and cloned into the BacMam vector^60^, in-frame with a PreScission cleavage site, followed by eGFP, FLAG-tag and 10× His-tag at the C-terminus. Baculovirus was generated according to manufacturer’s protocol and amplified to P3. For protein expression, HEK293S GnTI^-^ cells (ATCC) was cultured in Freestyle 293 media (Life Technologies) supplemented with 2% (v/v) FBS (Gibco) and 0.5% (v/v) PenStrep (Gibco). Cells were infected with 10% (v/v) P3 baculovirus at 2.5-3×10^6^ ml^-1^ density. After 20 hours shaking incubation at 37°C in the presence of 8% CO_2_, 10 mM sodium butyrate (Sigma-Aldrich) was added to the cell culture and the incubation temperature was lowered to 30°C to boost protein expression. After 44-48 hours, the cells were harvested by centrifugation at 550 × g, and was subsequently resuspended with lysis buffer (20 mM Tris pH8, 150 mM NaCl, 10 μg mL^-1^ leupeptin, 10 μg mL^-1^ pepstatin, 1 μg mL^-1^ aprotinin, 1 mM phenylmethylsulphonyl fluoride (PMSF, Sigma). The cells were lysed by probe sonication (45 pulses, 3 cycles). The membranes were subsequently solubilized by addition of 40mM DDM and 4mM CHS, followed by gentle agitation at 4°C for 1 hour. The solubilized lysate was cleared by centrifugation at 16,000 × g for 30 min to remove insoluble material. The supernatant was subsequently incubated with anti-FLAG M2 resin (Sigma-Aldrich) at 4°C for 1 hour. The resin was then packed onto a gravity-flow column, and washed with 10 column volumes of high-salt wash buffer (20 mM Tris pH 8, 300 mM NaCl, 5mM ATP, 10mM MgSO_4_, 0.07% digitonin), followed by 10 column volumes of wash buffer (20 mM Tris pH 8, 150 mM NaCl, 0.07% digitonin). Protein was then eluted with 5 column volumes of elution buffer (20 mM Tris pH 8, 150 mM NaCl, 0.07% digitonin, 100 μg mL^-1^ FLAG peptide). The eluted protein was concentrated with a 100kDa-cutoff spin concentrator (Millipore), after which 1:10 (w/w) PreScission protease was added to the eluted protein and incubated at 4°C for 1 h to cleave C-terminal tags. The mixture was further purified by injecting onto a Superdex 200 Increase (Cytiva) size-exclusion column equilibrated with GF buffer (20 mM Tris pH 8, 150 mM NaCl, 0.07% digitonin). The peak fractions were pooled and concentrated for cryo-EM sample preparation.

### Cryo-EM sample preparation

The peak fractions from final size exclusion chromatography were concentrated to 4-8 mg ml^-1^. For apo-OCT1_CS_ sample, a final concentration of 2% DMSO was added. For VPM-OCT1_CS_, 1 mM Verapamil (Sigma) was added to the sample approximately 30 minutes prior to vitrification. For DPH-OCT1_CS_ sample, 1mM diphenhydramine (Sigma) was added to the protein sample ∼30 minutes prior to vitrification. For MPP^+^-OCT2_CS_ sample, 1 mM MPP^+^ iodide (Sigma) was added to the protein sample ∼45 minutes prior to vitrification. All liganded samples maintain a 2% (v/v) DMSO concentration. Using Leica EM GP2 Plunge Freezer operated at 4°C and 95% humidity, 3 µL sample was applied to a freshly glow-discharged UltrAuFoil R1.2/1.3 300 mesh grids (Quantifoil), blotted with Whatman No. 1 filter paper for 1-1.5 seconds then plunge-frozen in liquid-ethane cooled by liquid nitrogen.

### Cryo-EM data collection

Apo-OCT1_CS_, DPH-OCT1_CS_ and MPP^+^-OCT2_CS_ datasets were collected using a Titan Krios (Thermo Fisher) transmission electron microscope operating at 300 kV equipped with a K3 (Gatan) detector in counting mode behind a BioQuantum GIF energy filter with slit width of 20eV, using Latitude S (Gatan) single particle data acquisition program. For Apo-OCT1_CS_ , DPH-OCT1_CS_, movies were collected at a nominal magnification of 81,000× with a pixel size of 1.08 Å/px at specimen level. Each movie contains 60 frames over 3.7 s exposure time, using a nominal dose rate of 20 e^-^/px/ּּּּּs, resulting a total accumulated dose of ∼60e^-^/Å^2^. For MPP^+^-OCT2_CS_, movies were collected at a nominal magnification of 81,000× with a pixel size of 1.08 Å/px at specimen level. Each movie contains 40 frames over 2.4 s exposure time, using a nominal dose rate of 30 e^-^/px/ּּּּּs, resulting a total accumulated dose of ∼60e^-^/Å^2^. The nominal defocus range was set from -0.8 to – 1.8 μm.

VPM-OCT1_CS_ dataset was collected using a Talos Arctica (Thermo Fisher) operating at 200kV equipped with a K3 (Gatan) detector operated in counting mode, using SerialEM^61^ automated data acquisition program with modifications to achieve high speed^62^. Movies were collected at a nominal magnification of 45,000× with a pixel size of 0.88 Å/px at specimen level. Each movie contains 60 frames over 2.7 s exposure time, using a dose rate of 14 e^-^/px/ּּּּּs, resulting in a total accumulated dose of ∼40 e^-^/Å^2^. The nominal defocus range was set from -0.6 to -1.2 μm.

### Cryo-EM data processing Apo-OCT1_CS_

Beam-induced motion correction and dose-weighing for a total of 5,993 movies were performed using MotionCor2^63^. Contrast transfer function parameters were estimated using Gctf^64^ or CTFFIND4^65^. Micrographs showing less than 6 Å estimated CTF resolution were discarded, leaving 5,304 micrographs. A subset of 100 micrographs were used for blob picking in cryoSPRAC^66, 67^, followed by 2D classification to generate templates for template-based particle picking. A total of 5.51 million particles were picked, followed by particle extraction with a 64- pixel box size at 4× binning (4.32 Å/pixel). A reference-free 2D classification was performed to remove obvious junk classes, resulting in a particle set of 1.52 million particles. Followed by 2D clean-up, an iterative ab initio triplicate procedure was performed in cryoSPARC, as described previously^58^. Specifically, three parallel ab initio reconstructions jobs were performed using identical settings (Initial resolution 35 Å, final resolution 12 Å, initial minibatch size 150, final minibatch size 600, class similarity 0, otherwise default settings were used). After the three parallel jobs conclude, particles from the class showing better protein features were selected from each job and combined, duplicates removed, then subjected to the next round of ab initio reconstruction triplicates with iteratively higher resolution limits. The same process was repeated multiple times until a reasonable reconstruction, showing acceptable protein features, was obtained. After 1 initial iteration in triplicate (initial resolution 20 Å, final resolution 10 Å), the remaining 1.34 million particles were re-extracted, re-centered using a 200 pixel box size, 2× binning (2.16 Å/pixel), resulting in a 100 pixel particle box size. The same iterative triplicate ab initio reconstruction procedure was performed for 11 iterations, with incrementally increasing initial/final resolutions (from 12 Å initial, 8 Å final, 4.5 Å final). The 11-iteration run was chosen because subsequent 12^th^ and 13^th^ iteration failed to improve map quality. A total of 458,246 particles were subsequently re-extracted and re-centered without binning with a 200 pixel box size (1.08 Å/pixel), followed by 7 rounds of ab initio reconstruction triplicates, resulting in 243,986 particles. A 3D volume (from earlier an *ab initio)* showing clear protein features was used as a projection template for a second round of particle picking. A 1.69 million particle set was picked using template-based picking in cryoSPARC, and a similar 2D-classification followed by iterative ab initio reconstruction triplicates as described before were performed, except that only 6 iterations were performed this time, as the particle set were significantly more homogenous. The resulting 415,943 particles were combined with the initial clean stack (243,986 particles), with duplicates removed for a total of 573,089 distinct particles. These particles were subjected to non-uniform (NU) refinement and local refinement, resulting in a 3.93 Å resolution reconstruction. To further improve map quality and resolution, the particle set was transferred to RELION 3.1^68^ and subjected to Bayesian polishing, followed by 3D classification without image alignment (K=8, T=40). One good class, containing 102,607 particles and exhibiting the best OCT1 features was selected and subjected to 3D refinement and Bayesian polishing. Polished particles were then imported to cryoSPARC and subjected to NU-refinement followed by local refinement, resulting in a 3.57 Å final reconstruction. Local resolution was estimated using cryoSPARC.

### DPH-OCT1_CS_

DPH-OCT1_CS_ dataset was processed similarly to that for Apo-OCT1_CS_ with minor modifications. Beam-induced motion correction and dose-weighing for a total of 7,145 movies were performed using MotionCor2^63^. Contrast transfer function parameters were estimated using Gctf ^64^or CTFFIND4^65^. Micrographs showing less than 4 Å estimated CTF resolution were discarded, leaving 2,233 micrographs. Template picking was performed in cryoSPRAC^66, 67^, using templates generated from a 3D-volume obtained from earlier processing attempts. A total of 693,720 particles were picked, followed by particle extraction with a 64-pixel box size at 4× binning (4.32 Å/pixel). A reference-free 2D classification was performed to remove obvious junk classes, resulting in a particle set of 638,957 particles. Followed by 2D clean-up, then iterative ab initio reconstruction triplicate runs were performed as described in the previous section. After 6 iterations with a progressively increasing resolution range, a 201,100 particle set was obtained, producing a 4.32 Å resolution reconstruction from NU-refinement. The particle set was subsequently imported to RELION^68^, and subjected to Bayesian polishing, followed by CTF refinement (beam tilt refinement only), followed by another Bayesian polishing job. The polished particles were transferred to cryoSPARC and subjected to two iterations of ab initio triplicates. The resulting 189,183 particles were subjected to NU-refinement and Local Refinement, producing the final map at 3.77 Å. Local resolution was estimated using cryoSPARC.

### VPM-OCT1_CS_

VPM-OCT1_CS_ dataset was processed similarly to that for Apo-OCT1_CS_ and DPH-OCT1_CS_ with minor modifications. Two datasets from two distinct grids, containing 4,050 and 2,057 movies, were subjected to beam-induced motion correction and dose-weighing in MotionCor2^63^. Contrast transfer function parameters were estimated using CTFFIND4^65^. Micrographs showing less than 4.0 Å estimated CTF resolution were discarded, leaving 3,249 micrographs. Template picking was performed in cryoSPARC^66, 67^, followed by particle extraction with a 64-pixel box size (4x binning, 3.52 Å/pixel) and 2D classification. A total of 495,998 particles corresponding to good 2D classes were selected, followed by particle extraction with a slightly larger box size (80-pixel box size at 4× binning; 3.52 Å/pixel). Following 2D clean-up and particle re-extraction, then iterative ab initio reconstruction triplicate runs were performed as described in the previous section. A total of 10 iterations were performed, followed by particle re-extraction with a 256-pixel box size (1x binning, 0.88 Å/pixel) and NU-refinement, producing 3.8 Å resolution reconstruction containing 89,771 particles. The stack was then transferred to RELION^68^ for two consecutive Bayesian polishing runs which helped boost resolution. The stack was then transferred back to cryoSPARC for final runs of NU refinement and Local Refinement, resulting in a 3.45 Å map. Local resolution was estimated using cryoSPARC.

### MPP^+^-OCT2_CS_

MPP^+^-OCT1_CS_ dataset was processed similarly to that for Apo-OCT1_CS_ with minor modifications. Beam-induced motion correction and dose-weighing for a total of 8,029 movies were performed using MotionCor2^63^. Contrast transfer function parameters were estimated using Gctf ^64^or CTFFIND4^65^. Micrographs showing less than 4 Å estimated CTF resolution were discarded, leaving 3,366 micrographs. Template-based picking was performed in cryoSPRAC^66, 67^, using templates projected from a 3D-volume obtained from earlier processing attempts on this dataset. A total of 1,141,906 particles were picked, followed by particle extraction with a 64-pixel box size at 4× binning (4.32 Å/pixel). A reference-free 2D classification was performed to remove obvious junk classes, resulting in a particle set of 1,044,268 particles. Following 2D clean-up, iterative ab initio reconstruction triplicate runs were performed as described in the previous section. After 25 iterations with a progressively increasing resolution range, a 92,217-particle set was obtained, producing a 4.09 Å resolution reconstruction from Local Refinement. The particle set was subsequently imported to RELION^68^, and subjected to two successive iterations of Bayesian polishing. The polished particles were transferred to cryoSPARC and subjected to four more iterations of ab initio triplicates. The resulting 73,474 particles were subjected to NU-refinement and Local Refinement, producing the final map at 3.61 Å resolution. Local resolution was estimated using cryoSPARC.

### Model Building and Refinement

All manual model building was performed in Coot^69^ with ideal geometry restraints. A *de novo* initial model was built to a 3D reconstruction of VPM-OCT1_CS_ cryo-EM map, followed by further manual model building and adjustment. Idealized CIF restraints for ligands were generated in eLBOW (in PHENIX software suite^70^) from isomeric SMILES strings. Further manual adjustments were performed on both protein and ligands after placement, to ensure correct stereochemistry and good geometries. The manually refined coordinates were subjected to phenix-real.space.refine in PHENIX with global minimization, local grid search and secondary structure restraints. MolProbity^71^ was used to help identify errors and problematic regions. The refined VPM-OCT1_CS_ cryo-EM structure was then rigid-body fit into the apo-OCT1_CS_, DPH-OCT1_CS_ maps, followed by manual coordinate adjustments, ligand placement and adjustments for DPH- OCT1_CS_, followed by phenix-real.space.refine in PHENIX. Moreover, the VPM-OCT1_CS_ was rigid-body fit into MPP^+^-OCT1_CS_ with proper adjustments to the sequence, followed by manual adjustments, ligand placement, followed by phenix-real.space.refine in PHENIX. The Fourier shell correlation (FSC) of the half- and full-maps against the model, calculated in Phenix, were in good agreement for all five structures, indicating that the models did not suffer from over-refinement (Extended Data Figure 3). Structural analysis and illustrations were performed using Open Source PyMOL, UCSF Chimera ^72^ and UCSF Chimera X^73^.

### Molecular Dynamics Simulations and docking

All-atom molecular dynamics (MD) simulations in explicit solvents and POPC bilayer membranes were performed using the cryo-EM structures of OCT1_CS_ in two holo states containing DPH (2 different conformations) and VPM (2 possible poses). The systems were assembled using the CHARMM-GUI web server^74–76^. Each system was solvated in TIP3P water and neutralized with Na+ and Cl- ions at 0.15 M^77^. Five independent replicates were simulated for each system. Non-bonded van der Waals interactions were truncated between 10 and 12 Å using a force-based switching method.^78^ The long-range electrostatic interactions were calculated using the Particle-mesh Ewald summation^79^. The systems were equilibrated following the CHARMM-GUI *Membrane Builder* protocol.^75, 76^ The production runs were performed in the NPT (constant particle number, pressure, and temperature) for 500 ns at 303.15 K and 1 bar with hydrogen mass repartitioning^80, 81^ using the CHARMM36m force field (protein and lipid) and CGenFF (diphenhydramine and verapamil)^82, 83^. All simulations were performed with the OpenMM package^84^.

Ligand binding stability was evaluated using ligand RMSD by superimposing the transmembrane coordinates of the protein structure throughout MD trajectory and calculating ligand RMSD using CHARMM.^85^ Probability density maps of the amine nitrogen atom were calculated using Chimera^72^.

A docking/MD simulation workflow to determine the ligand binding pose of 10 ligands in the binding pocket of hOCT1 was performed following the CHARMM-GUI *High-Throughput Simulator* protocol (Figure 4a).^50, 51^ Two structures were used, OCT1_CS_-DPH and OCT1_CS_-VPM. Rigid docking was conducted using AutoDock Vina^86^. The center-of-mass coordinate of the bound ligand (DPH & VPM) were used to determine the docking search space. Ligand docking was performed on a cubic box search space with 22.5 Å edges. For each ligand, top 10 binding poses based on docking scores were collected for subsequent high-throughput MD simulations and rescoring using MD ligand RMSD & molecular mechanics with the Poisson-Boltzmann electrostatic continuum solvation and surface area (MMPBSA)^87^. MD simulation systems were built similarly to the protocol described above. The production runs were performed for 50 ns for each protein-ligand complex structure. Ligand RMSD was used to determine the ligand binding stability of each binding pose. Only binding poses with < 3 Å average ligand RMSD were selected as good binding modes. Subsequently, molecular mechanics with MMPBSA calculations were performed on 51 protein-ligand structures to determine the best binding pose for each ligand. MMPBSA calculations were done following the protocol previously described ^51^.

## Acknowledgements

Cryo-EM data were screened and collected at the Duke University Shared Materials Instrumentation Facility (SMIF), the UNC CryoEM core facility, and the National Institute of Environmental Health Sciences (NIEHS). We thank Nilakshee Bhattacharya at SMIF, and Jared Peck of the UNC CryoEM Core Facility for assistance with the microscope operation. This research was supported by a National Institutes of Health (R01GM137421 to S.-Y.L and R01GM138472 to W.I.), the National Institute of Health Intramural Research Program; US National Institutes of Environmental Health Science (ZIC ES103326 to M.J.B.) and a National Science Foundation grant MCB-1810695 (W.I.). DUKE SMIF is affiliated with the North Carolina Research Triangle Nanotechnology Network, which is in part supported by the NSF (ECCS- 2025064). The UNC CryoEM core facility is supported by NIH grant P30CA016086.

## Author Contributions

Y.S. conducted biochemical preparation, sample freezing, grid screening, data collection, data processing and single particle 3D reconstruction as well as surface expression experiments, N.J.W. performed radiotracer uptake assays, data processing and single particle 3D reconstruction, all under the guidance of S.-Y.L. J.G.F. performed part of radiotracer and surface expression experiments. N.J.W. Y.S. and S.-Y.L. performed model building and refinement. H.G. carried out all MD simulations as well as docking studies under the guidance of W.I. K.J.B. helped with part of cryo-EM sample screening and provide advice on sample freezing under the guidance of M.J.B. N.J.W. Y.S. and S.-Y.L. wrote the paper.

## Competing Interests

The authors declare no competing interests.

## Data Availability

Atomic coordinates have been deposited in the Protein Data Bank with the PDB IDs 8ET6 (Apo- OCT1_CS_), 8ET7 (DPH-OCT1_CS_), and 8ET8 (VPM-OCT1_CS_), 8ET9 (MPP^+^-OCT2_CS_), respectively. The reconstructed cryo-EM maps have been deposited in the Electron Microscopy Data Bank with the IDs EMD-28586 (Apo-OCT1_CS_), EMD-28587 (DPH-OCT1_CS_), and EMD-28588 (VPM- OCT1_CS_), EMD-28589 (MPP^+^-OCT2_CS_), respectively.

## Extended Data Figures

**Extended Data Figure 1.**
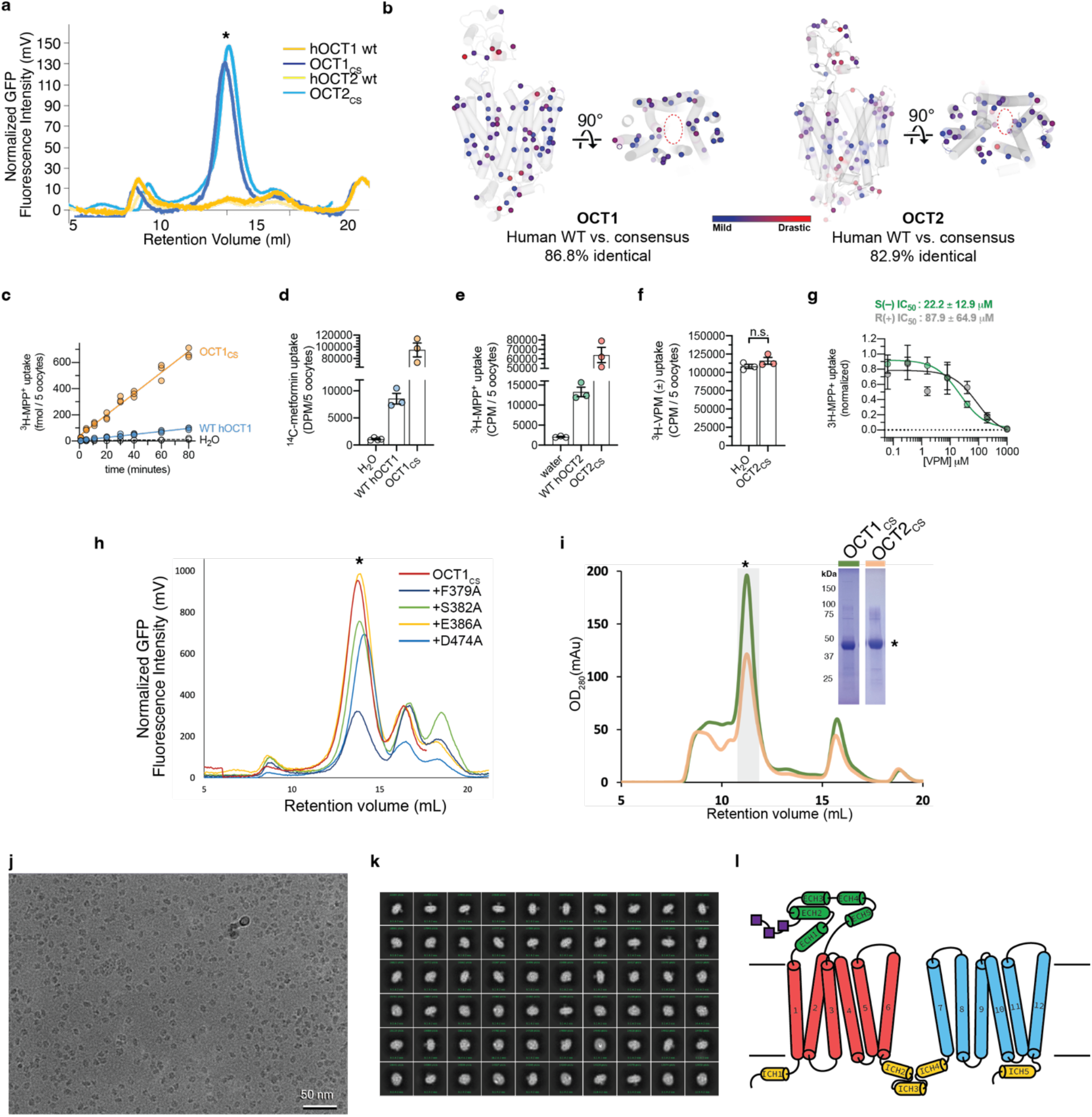
Consensus mutagenesis, protein biochemistry, and cryo-EM analysis of OCT1_CS_. **a,** FSEC traces showing strong monodisperse peak of OCT1_CS_-GFP, WT hOCT1-GFP, OCT1_CS_- GFP and WT hOCT2-GFP which exhibits no discernable peak corresponding to target protein (expression performed in HEK293T cells). **b,** Map of all residues in OCT1_CS_ and OCT2_CS_ that deviate from WT hOCT1. The residues are colored based on their conservation score from MAFFT alignment. Blue spheres indicate mildly changed, while red spheres indicate drastic changes. Only two residues differ from hOCT1 (Y36 and F446 in OCT1_CS_ and C36 and I446 in hOCT1) in the central ligand binding cavity. **c,** Time-dependent accumulation of 10 nM [^3^H]-MPP+ in WT hOCT1 and OCT1_CS_ expressing oocytes (n = 3 per timepoint). **d,** Raw uptake values for controls in the OCT1 [^14^C]-metformin uptake experiments, corresponding to Fig. 1e. **e,** Raw uptake values for controls in the OCT2 [^3^H]-MPP+ uptake experiments, corresponding to Fig. 1f. **f,** [^3^H]- verapamil (100 nM) does not accumulate in OCT2_CS_-injected oocytes over the course of 30 min (n =3, individual values and mean ± S.E.M), demonstrating it is not an OCT2_CS_ substrate. **g,** Stereoselectivity of verapamil inhibition against OCT2_CS_ mediated [^3^H]-MPP+ uptake (100 nM radiotracer, 30 minute uptake, n =3, mean ± S.E.M) **h,** FSEC traces showing expression of selected OCT1_CS_ mutants in HEK293T cells. **i,** Representative size-exclusion chromatography trace (left) and SDS-PAGE (right) of purified OCT1_CS_ and samples OCT1_CS_ used for cryo-EM grid preparation. **j,** Representative micrograph of a OCT1_CS_ sample. OCT2 _CS_ behaves similarly under cryo-EM with OCT1 _CS_. **k,** Representative 2D classes from a OCT1_CS_ dataset. **l**, Secondary structure topology of OCT1 and OCT2.

**Extended Data Figure 2.**
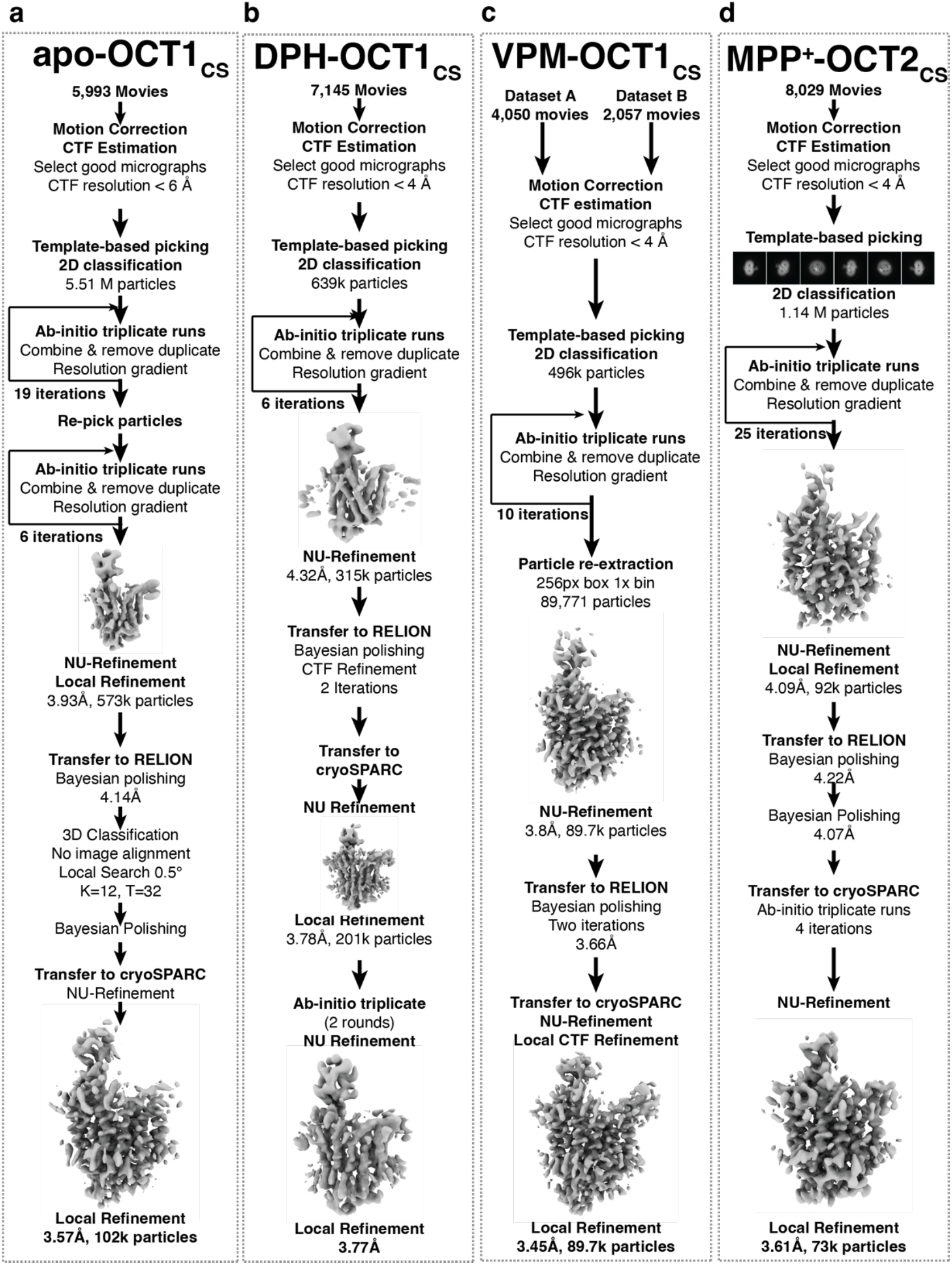
Cryo-EM data processing workflow. **a-e,** cryo-EM data processing workflow for apo-OCT1_CS_, DPH-OCT1_CS_, VPM-OCT1_CS_, and MPP^+^-OCT2_CS_ datasets, respectively.

**Extended Data Figure 3.**
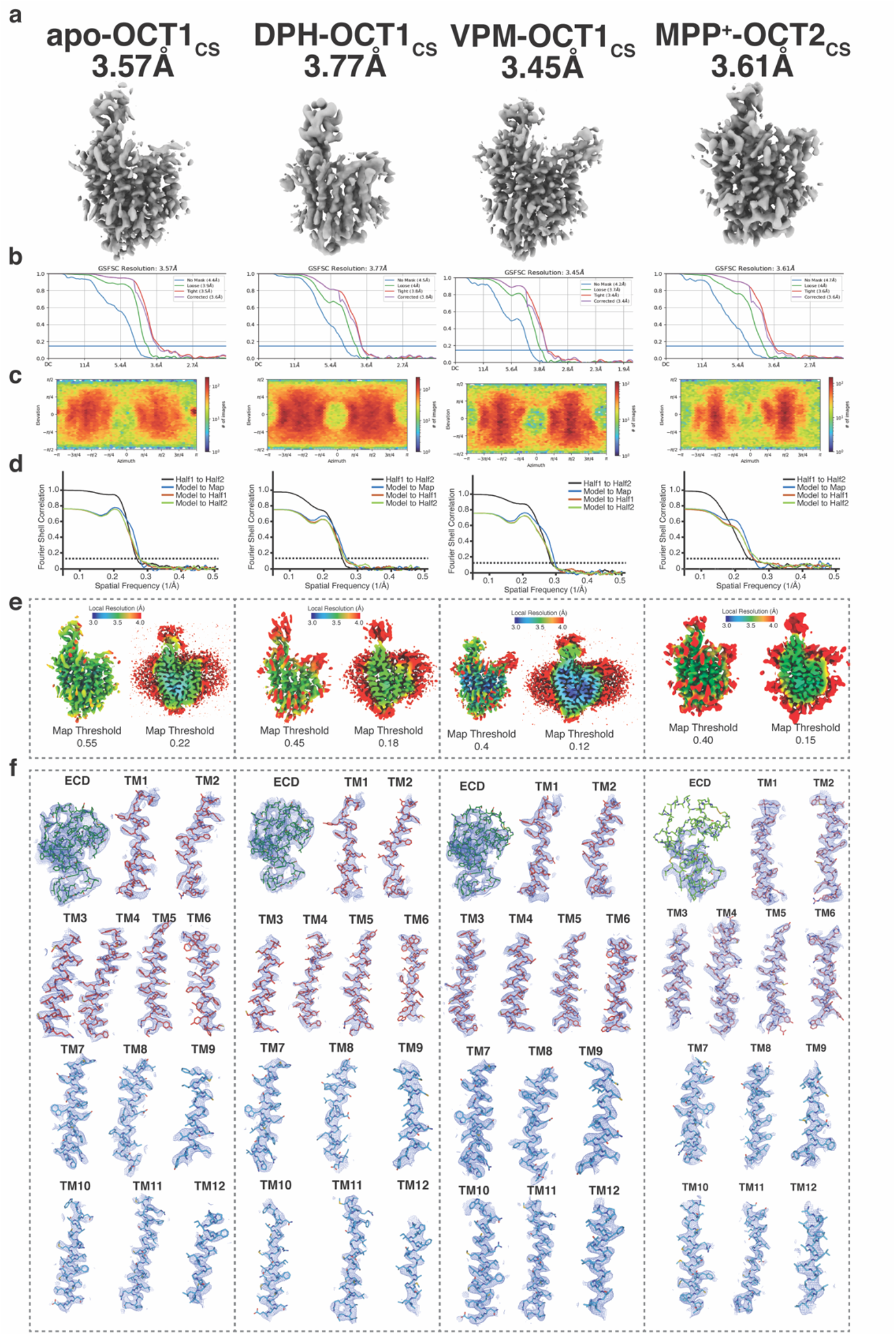
Cryo-EM data validation. **a,** Final cryo-EM reconstructions. **b,** Fourier-shell correlation for the final reconstruction, generated from cryoSPARC. **c,** projection orientation distribution map for the final reconstruction, generated from cryoSPARC. **d,** Map-to-model correlation plots. **e,** Local Resolution plots. **f,** cryo- EM maps for secondary structure segments. From left to right are cryo-EM data validations for apo-OCT1_CS_, DPH-OCT1_CS_, VPM-OCT1_CS_, and MPP^+^-OCT2_CS_ datasets, respectively.

**Extended Data Figure 4.**
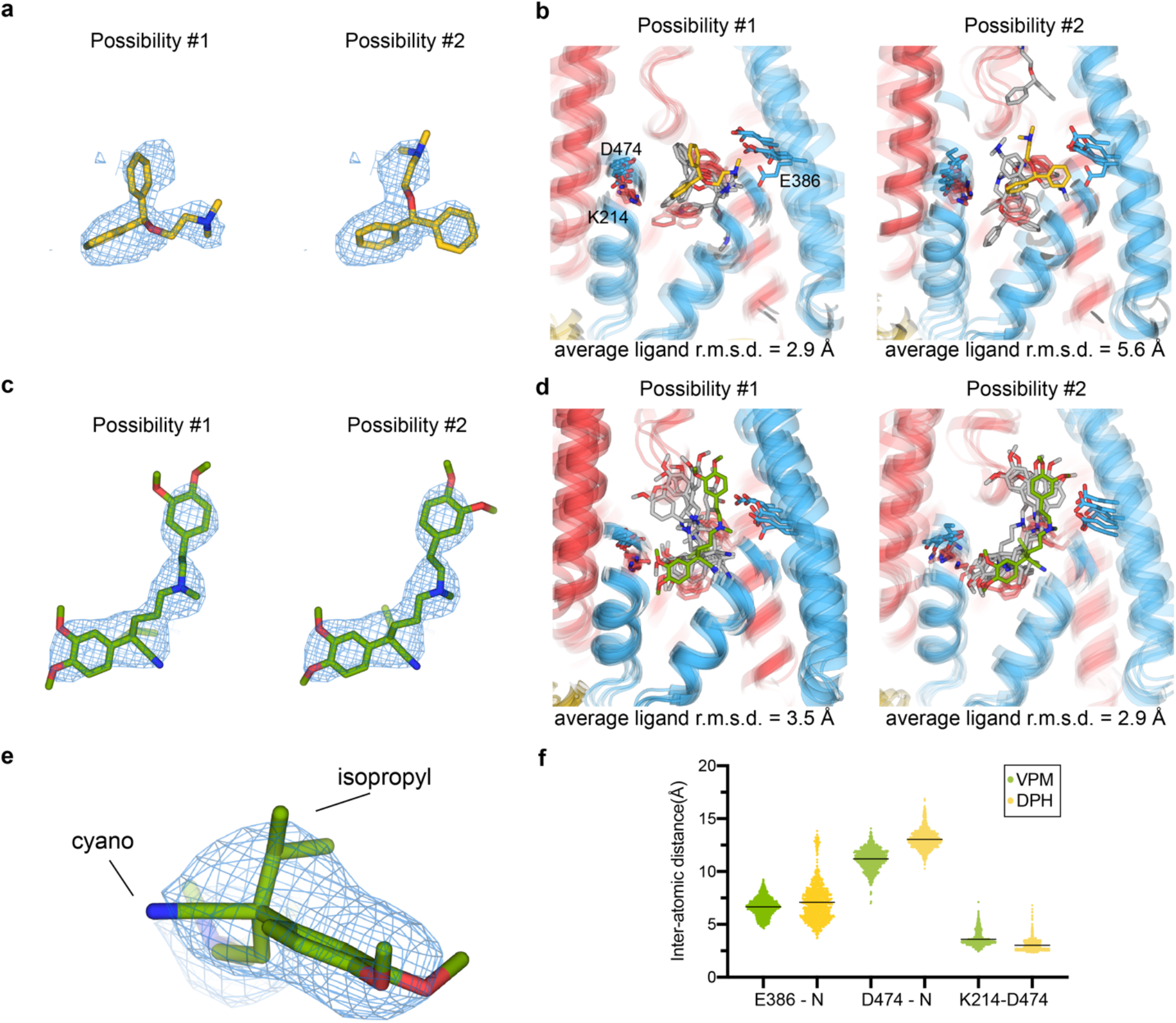
Validation of ligand binding poses with molecular dynamics simulations. **a**, Two possible poses for DPH molecule placement based on the cryo-EM reconstruction. **b**, Final MD frame for 5 replicas of DPH-OCT1_CS_ MD simulations (500ns) for the two proposed poses, where possibility #1 is more stably bound at the site. **c,** Two possible poses for S(–)-VPM based on the cryo-EM reconstruction. **d,** Final MD frame for 5 replicas of VPM-OCT1_CS_ MD simulations (500ns), for the two proposed poses, where possibility #2 is more stable. **e,** Zoom-in view of the cryo-EM map and model of the VPM chiral center. **f,** Inter-atomic distances between select chemical groups during the MD-simulations of drug-bound OCT1_CS_ (scatter plot showing individual values extracted per MD frame, compiled from all 5 replicas per condition).

**Extended Data Figure 5.**
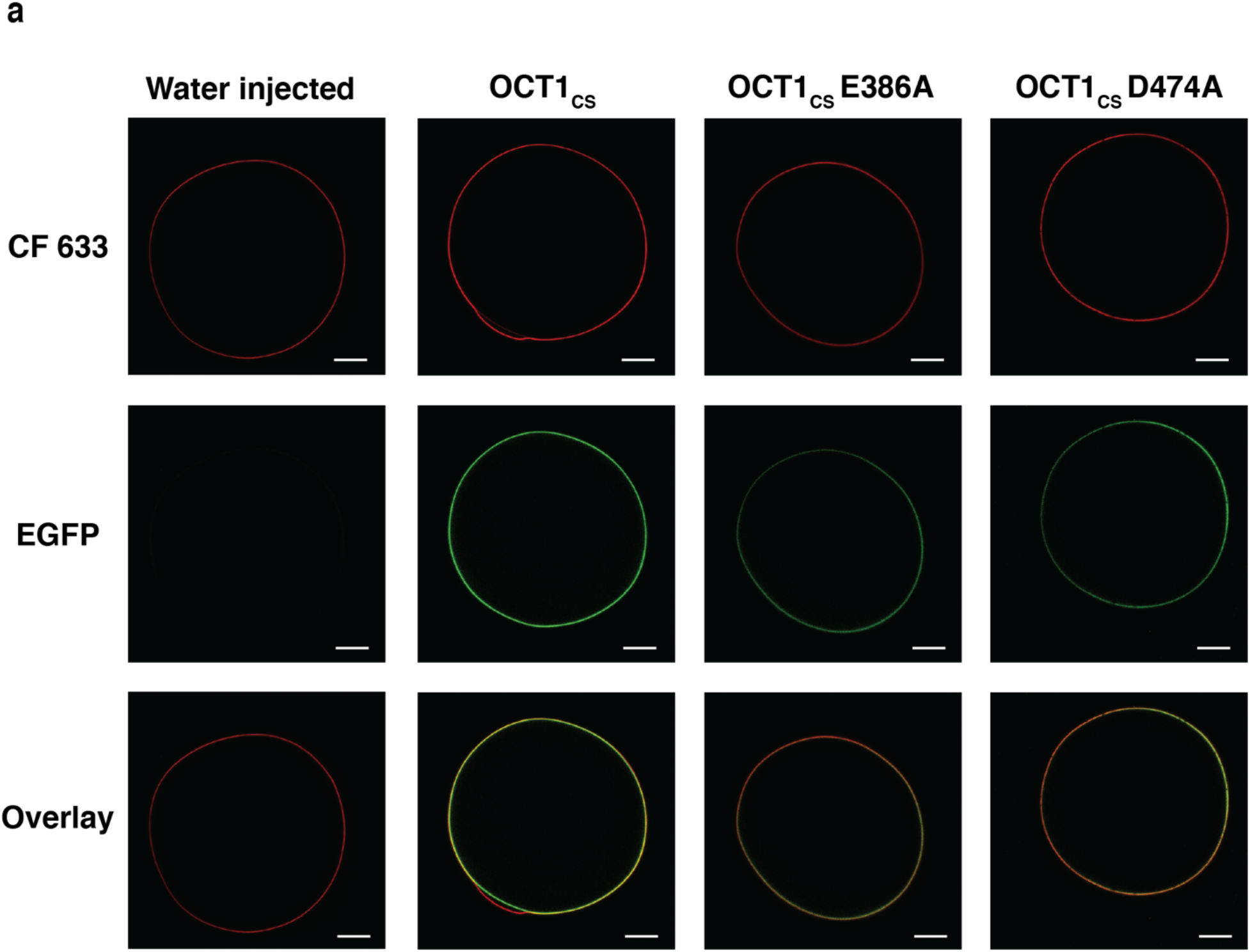
Surface expression of hOCT1-WT, OCT1_CS_ and mutants. Representative confocal microscopy images showing surface expression of OCT1_CS_ and relevant mutants in *Xenopus laevis* oocytes used for radiotracer uptake studies. OCT1_CS_-GFP +E386A and +D474A shows expression level slightly lower than that in OCT1_CS_. Scale bars represent 200 μm. Similar results were observed in 6-8 additional biological replicates per condition.

**Extended Data Figure 6.**
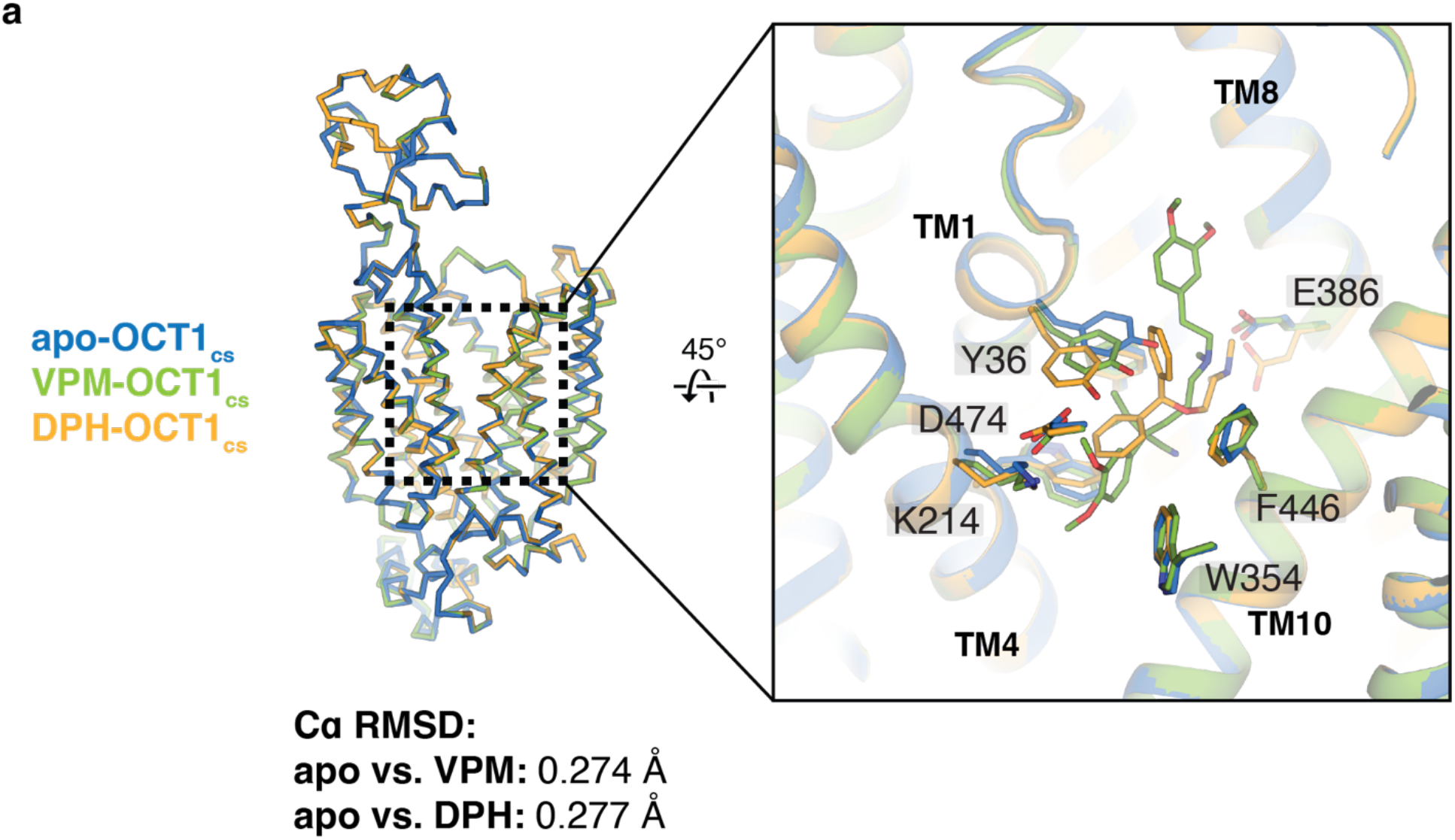
Ligand-induced local conformational changes in OCT1_CS_. **a**, Structural overlay of apo-OCT1_CS_ (marine), VPM-OCT1_CS_ (green) and DPH-OCT1_CS_ (yellow), showing that no large conformational changes are present among the three structures. While other residues remain relatively stable, Y36 exhibits considerable rotamer movement among the three structures.

**Extended Data Figure 7.**
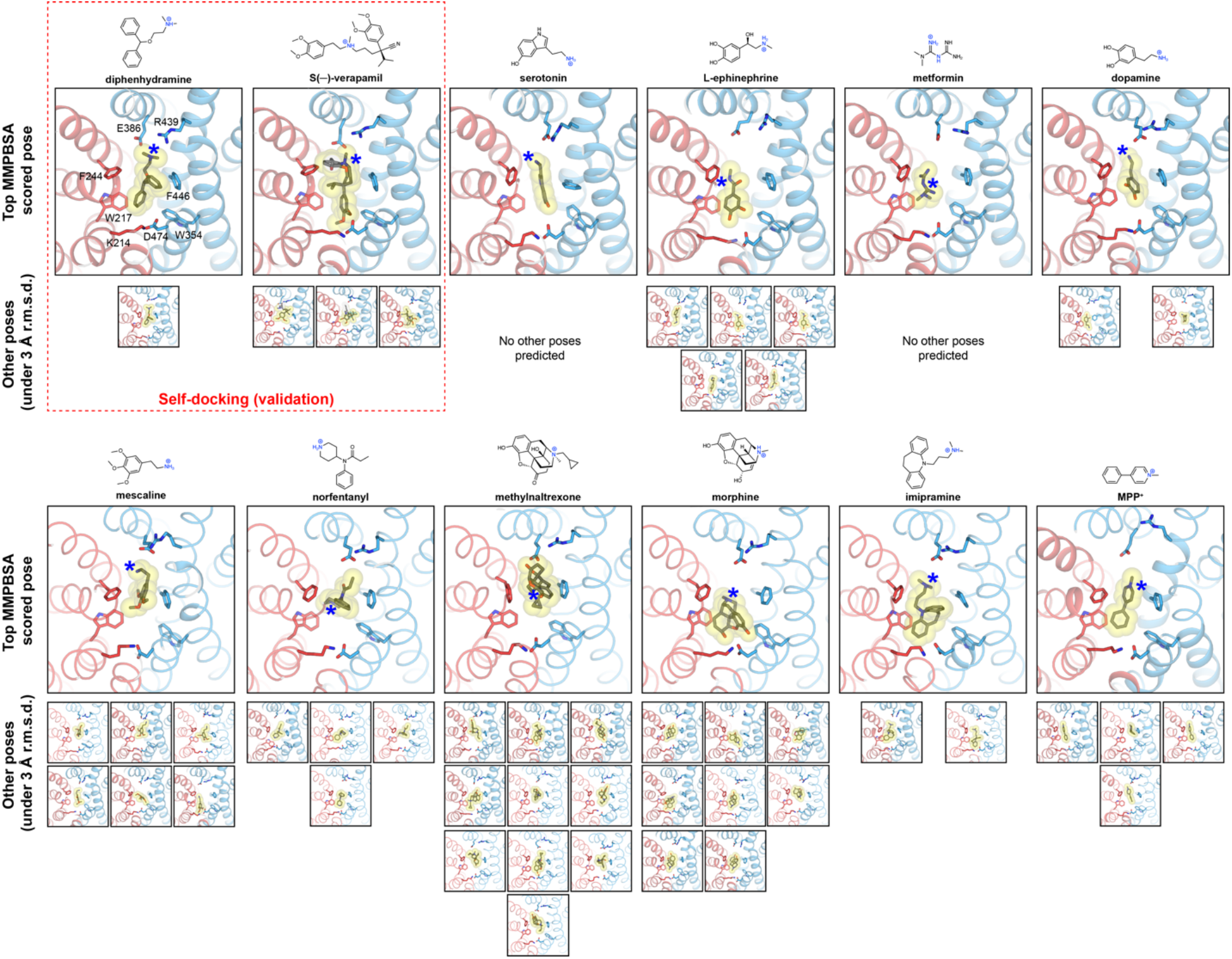
*In silico* ligand docking. In-silico docking and short-time scale (50ns) MD simulations for serotonin, epinephrine, metformin, dopamine, mescaline, norfentanyl, methylnaltrexone, morphine, imipramine and MPP^+^, respectively. For each ligand, Top MMPBSA scored poses are shown in the large panels, with other candidate poses (under 3 Å ligand r.m.s.d. at the conclusion of the simulation) shown below. Self-docking runs of DPH and VPM shown at top left for validation.

**Extended Data Fig. 8.**
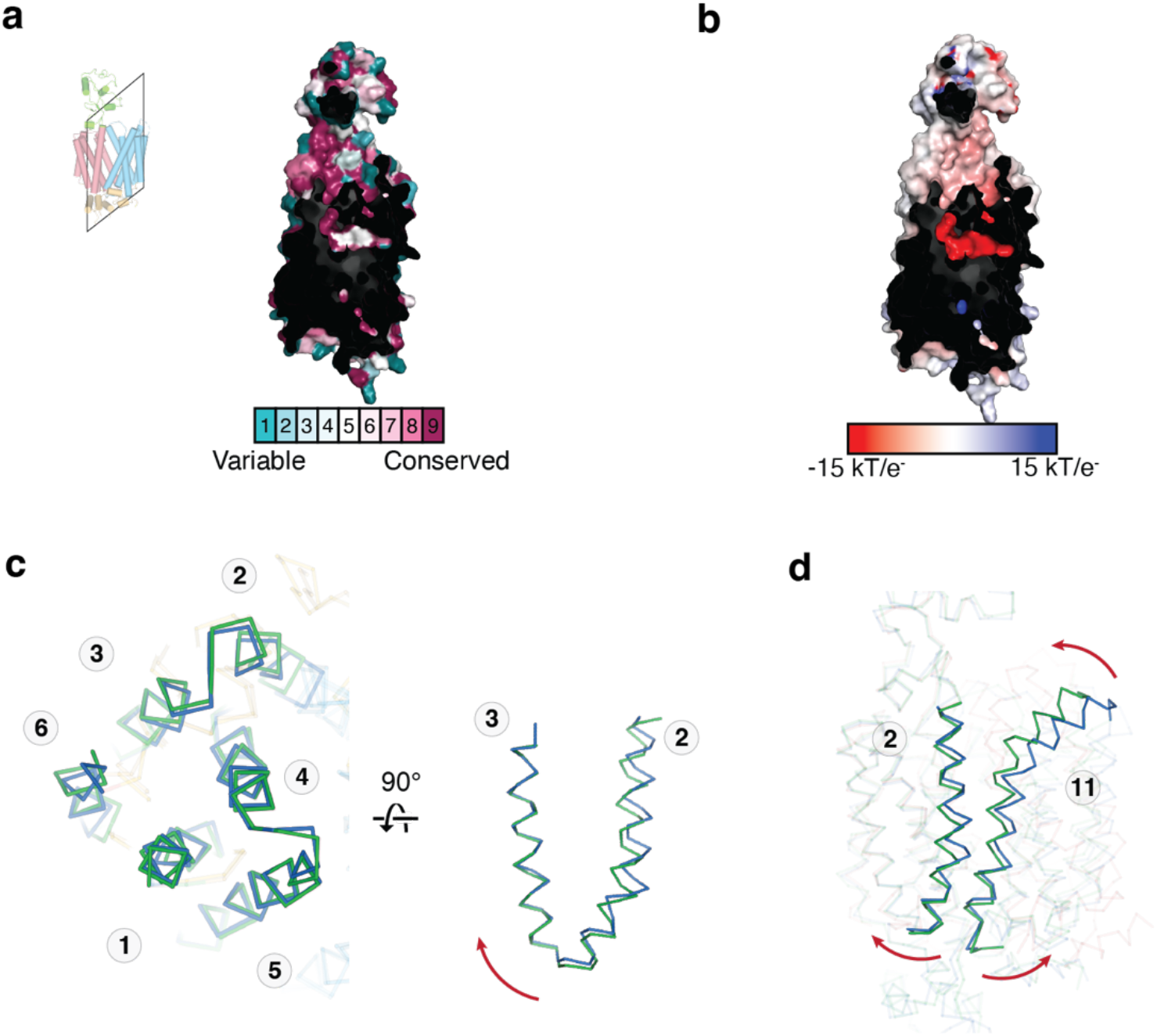
Local conformational changes associated with OCT gating. **a,** ConSurf plot for OCT2cs and OCT2 homologs. **b,** Electrostatics surface of outward occluded OCT2, calculated by APBS. **c,** local conformational changes in the N-lobe from outward open (blue) to outward occluded (green) conformations. **d,** Concerted local conformational changes in TM2 and 11 leads to extracellular gate formation.

**Supplemental Figure 1.**
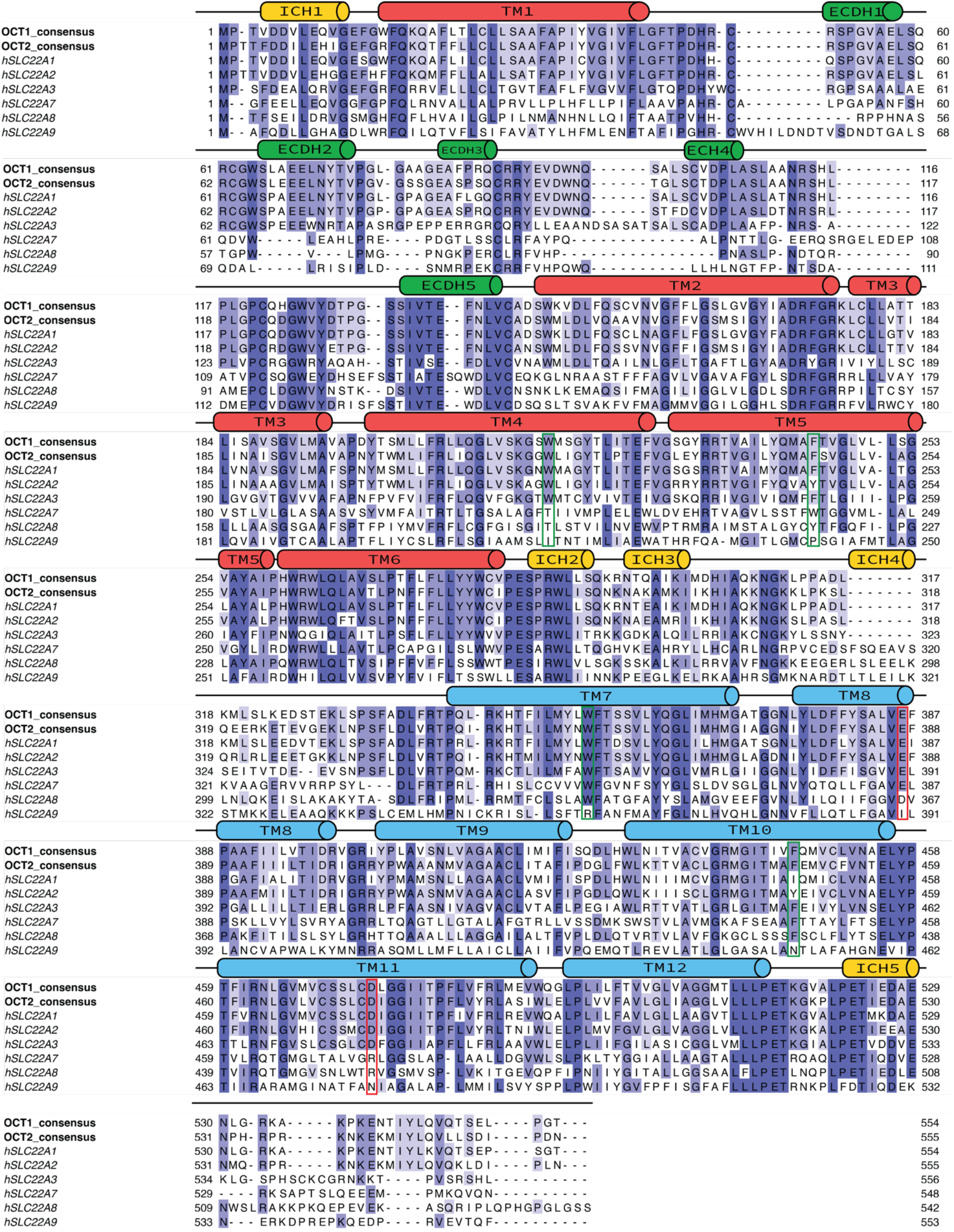
Multiple Sequence alignment. Multiple sequence alignment of OCT1_CS_, human OCTs (SLC22A1-3), and representative human OATs (SLC22A7-9). Sequences are aligned using MAFFT^56^. E386 and D474 (numbering according to hOCT1) positions are highlighted in red and 217, 244, 354, and 446 (numbering according to hOCT1) in green.

## Extended Data Tables

**Extended Data Table 1.** Cryo-EM data collection, refinement, and validation statistics.

|  | Apo-OCT1<sub>CS</sub><br>(EMD-28586)<br>(PDB 8ET6) | DPH-OCT1<sub>CS</sub><br>(EMD-28587)<br>(PDB 8ET7) | VPM-OCT1<sub>CS</sub><br>(EMD-28588)<br>(PDB 8ET8) | MPP<sup>+</sup>-OCT2<sub>CS</sub><br>(EMD-28589)<br>(PDB 8ET9) |
| --- | --- | --- | --- | --- |
| **Data collection and processing** |  |  |  |  |
| Magnification | 81,000 | 81,000 | 45,000 | 81,000 |
| Voltage (kV) | 300 | 300 | 200 | 300 |
| Electron exposure (e<sup>-</sup>/Å<sup>2</sup>) | 60 | 60 | 40 | 60 |
| Defocus range (μm) | -0.8 to -1.8 | -0.8 to -1.8 | -0.6 to -1.6 | -0.8 to -2.0 |
| Pixel size (Å) | 1.08 | 1.08 | 0.88 | 0.5347 |
| Symmetry imposed | C1 | C1 | C1 | C1 |
| Initial particle images (no.) | 5,515,896 | 5,495,544 | 1,794,057 | 1,141,906 |
| Final particle images (no.) | 102,580 | 189,183 | 89,771 | 73,474 |
| Map resolution (Å) | 3.57 | 3.77 | 3.45 | 3.61 |
| FSC threshold | 0.143 | 0.143 | 0.143 | 0.143 |
| **Refinement** |  |  |  |  |
| Initial model used (PDB code) | VPM-OCT1<sub>CS</sub> | VPM-OCT1<sub>CS</sub> | - | VPM-OCT1<sub>CS</sub> |
| Map sharpening *B* factor (Å<sup>2</sup>) | -152.3 | -164.4 | -121.1 | -141.9 |
| Model composition |  |  |  |  |
| Non-hydrogen atoms | 8,117 | 8,157 | 8,188 | 7,697 |
| Protein residues | 532 | 532 | 532 | 517 |
| Ligands | NAG:2 | DPH:1, NAG:2 | VPM:1, NAG:2 | MPP:1 |
| *B* factors (Å<sup>2</sup>) |  |  |  |  |
| Protein | 67.72 | 84.59 | 59.60 | 113.01 |
| Ligand | 132.52 | 111.90 | 81.89 | 93.91 |
| R.m.s. deviations |  |  |  |  |
| Bond lengths (Å) | 0.003 | 0.002 | 0.003 | 0.005 |
| Bond angles (°) | 0.521 | 0.468 | 0.569 | 0.610 |
| Validation |  |  |  |  |
| MolProbity score | 1.49 | 1.38 | 1.47 | 1.46 |
| Clashscore | 4.90 | 3.91 | 4.39 | 8.46 |
| Poor rotamers (%) | 0.00 | 0.00 | 0.00 | 0.52 |
| Ramachandran plot |  |  |  |  |
| Favored (%) | 96.42 | 96.79 | 96.23 | 98.24 |
| Allowed (%) | 3.58 | 3.21 | 3.77 | 1.76 |
| Disallowed (%) | 0.00 | 0.00 | 0.00 | 0.00 |

**Extended Data Table 2.**
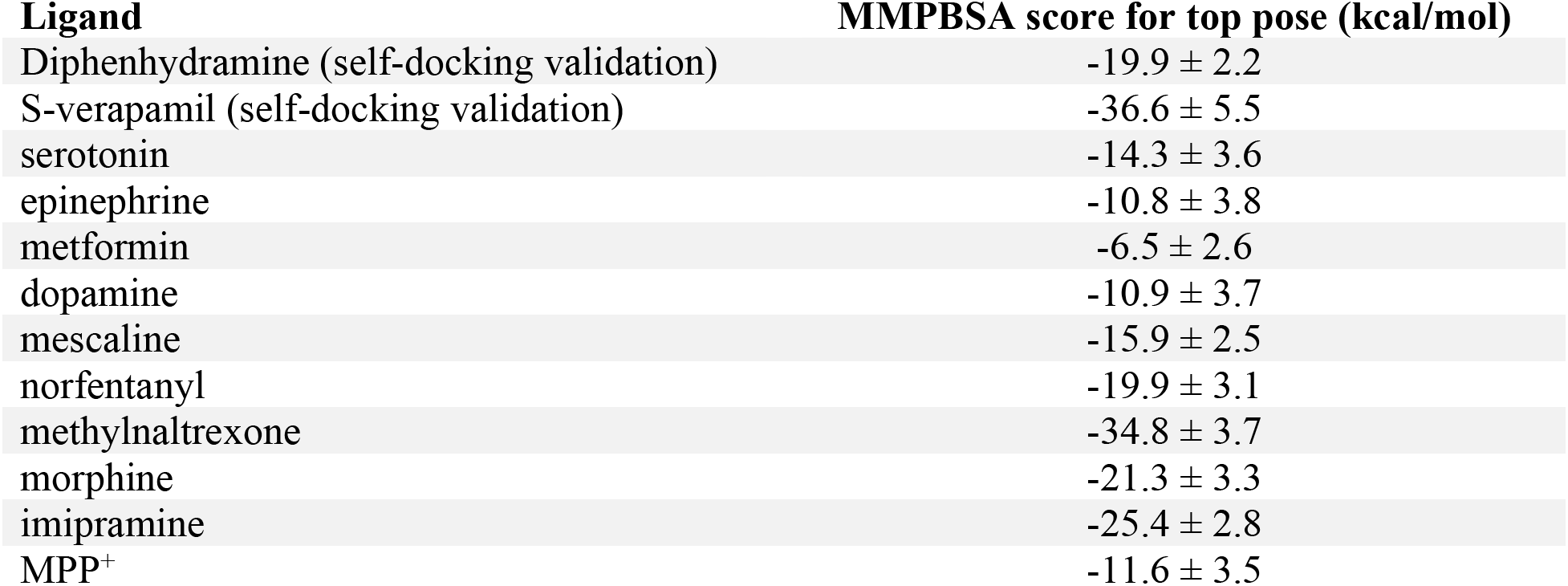
MMPBSA scores for top *in silico* docking poses. MMBPSA scores (kcal/mol) are shown as mean ± s.d.

| **Ligand** | **MMPBSA score for top pose (kcal/mol)** |
| --- | --- |
| Diphenhydramine (self-docking validation) | -19.9 ± 2.2 |
| S-verapamil (self-docking validation) | -36.6 ± 5.5 |
| serotonin | -14.3 ± 3.6 |
| epinephrine | -10.8 ± 3.8 |
| metformin | -6.5 ± 2.6 |
| dopamine | -10.9 ± 3.7 |
| mescaline | -15.9 ± 2.5 |
| norfentanyl | -19.9 ± 3.1 |
| methylnaltrexone | -34.8 ± 3.7 |
| morphine | -21.3 ± 3.3 |
| imipramine | -25.4 ± 2.8 |
| MPP<sup>+</sup> | -11.6 ± 3.5 |

## Notes

### Competing Interest Statement

The authors have declared no competing interest.

